# Soil conditions drive belowground trait space in temperate agricultural grasslands

**DOI:** 10.1101/2021.07.07.450881

**Authors:** Tom Lachaise, Joana Bergmann, Norbert Hölzel, Valentin H. Klaus, Till Kleinebecker, Matthias C. Rillig, Mark van Kleunen

**Affiliations:** Ecology, Department of Biology, University of Konstanz, Universitätsstrasse 10, D-78457 Konstanz, Germany; Institute of Biology, Freie Universität Berlin, Altensteinstr. 6, D-14195 Berlin, Germany; Leibniz Centre for Agricultural Landscape Research (ZALF), D-15374, Müncheberg, Germany; Berlin-Brandenburg Institute of Advanced Biodiversity Research (BBIB), D-14195 Berlin, Germany; Institute of Landscape Ecology, University of Münster, Heisenbergstr. 2, 48149 Münster; Institute of Agricultural Sciences, ETH Zürich, Universitätstr. 2, 8092 Zürich, Switzerland; Institute of Landscape Ecology and Resources Management, University of Giessen, Heinrich-Buff-Ring 26-32, D-35392 Giessen, Germany; Zhejiang Provincial Key Laboratory of Plant Evolutionary Ecology and Conservation, Taizhou University, Taizhou 318000, China

**Keywords:** clonal traits, environmental filtering, land-use, mycorrhiza, nitrogen, phosphorus, plant economics spectrum, root traits

## Abstract

1. Plant belowground organs perform essential functions, including water and nutrient uptake, anchorage, vegetative reproduction and recruitment of mutualistic soil microbiota. Determining how belowground traits jointly determine dimensions of the trait space and how these dimensions are linked to environmental conditions would further advance our understanding of plant functioning and community assembly.
2. Here, we investigated belowground plant-trait dimensionality and its variation along 10 soil and land-use parameters in 150 temperate grasslands plots. We used eight belowground traits collected in greenhouse and common garden experiments, as well as bud-bank size and specific leaf area from databases, for a total of 313 species, to calculate community weighted means (CWMs).
3. Using PCA, we found that about 55% of variance in CWMs was explained by two main dimensions, corresponding to a mycorrhizal ‘collaboration’ and a resource ‘conservation’ gradient. Frequently overlooked traits such as rooting depth, bud-bank size and root branching intensity were largely integrated in this bidimensional trait space. The two plant-strategy gradients were partially dependent on each other, with ‘outsourcing’ communities along the collaboration gradient being more often ‘slow’. These ‘outsourcing’ communities were also more often deep-rooting, and associated with soil parameters, such as low moisture and sand content, high topsoil pH, high C:N and low δ15N. ‘Slow’ communities had large bud-banks and were associated with low land-use intensity, high topsoil pH, and low nitrate but high ammonium concentrations in the soil. We did not find a substantial role of phosphorus-availability as an indicator along the ‘collaboration’ gradient.
4. In conclusion, the ‘collaboration’ and ‘conservation’ gradients previously identified at the species level scale up to community level in grasslands, encompass more traits than previously described, and vary with the environment.

## Introduction

Plant traits are of major interest as they determine plant functioning (Solbrig, 1993), covary with environmental conditions (Garnier, Navas, & Grigulis, 2016), and influence ecosystem functions (de Bello et al., 2010; Hanisch, Schweiger, Cord, Volk, & Knapp, 2020). Nevertheless, traits may have low predictive power (Klimešová, Tackenberg, & Herben, 2016; van der Plas et al., 2020), because there is limited understanding which and how many traits are needed in ecological studies (Shipley et al., 2016). An important step forward has been the grouping of multiple traits into a limited number of syndromes, with continuous variation in the form of gradients of plant strategies (Westoby, Falster, Moles, Vesk, & Wright, 2002; Wright et al., 2004; Chave et al., 2009; Pierce, Brusa, Vagge, & Cerabolini, 2013; Díaz et al., 2016; Klimešová, Martínková, & Herben, 2018; Bergmann et al., 2020; Roddy et al., 2020). For example, Diaz et al. (2016) showed that the variation in aboveground traits can be captured by a ‘size’ gradient representing the size of whole plants and plant organs, and an ‘economic’ gradient representing the leaf economics spectrum. A similar attempt has recently addressed root traits that identified a ‘conservation’ gradient and a ‘collaboration’ gradient as two independent axes of belowground plant economy (Weemstra et al. 2016; Kramer-Walter et al. 2016; Bergmann et al. 2020).

Bergmann et al. (2020) suggested that in the root economic space the ‘conservation’ gradient, ranging from ‘slow’ to ‘fast’, is related to carbon conservation and determined by root-tissue density and nitrogen content. In contrast, the ‘collaboration’ gradient, ranging from ‘do-it-yourself’ to ‘outsourcing’ of resource uptake to fungal partners, is reflected by specific root length and root diameter along with mycorrhizal colonization (Fig. 1a). Despite the recent progress in the understanding of trait dimensionality, several root traits with a high potential importance for plant functioning (Laliberté, 2017) were so far not integrated into the existing framework. For example, high root-branching intensity can be seen as an alternative to the reliance on mycorrhiza, and may be associated with specific root length for better local soil exploitation (Kong et al., 2014; Freschet et al., 2020; Ding et al., 2020). Thus it may be indicative of a ‘do-it-yourself’ strategy. Furthermore, rooting depth is also likely to be an important source of interspecific variation as it varies considerably across biomes (Schenk & Jackson, 2002), and could explain overyielding in grasslands via the mechanism of resource partitioning (Mommer et al., 2010; Mueller, Tilman, Fornara, & Hobbie, 2013). As deep-rooting species are able to take up nutrients and water from deeper soil layers, rooting depth might be part of the ‘fast’ strategy of the conservation gradient (Fig. 1a). However, up to now many of these belowground traits received much less attention, limiting a comprehensive understanding of the plant-soil interface.

**Figure 1.**
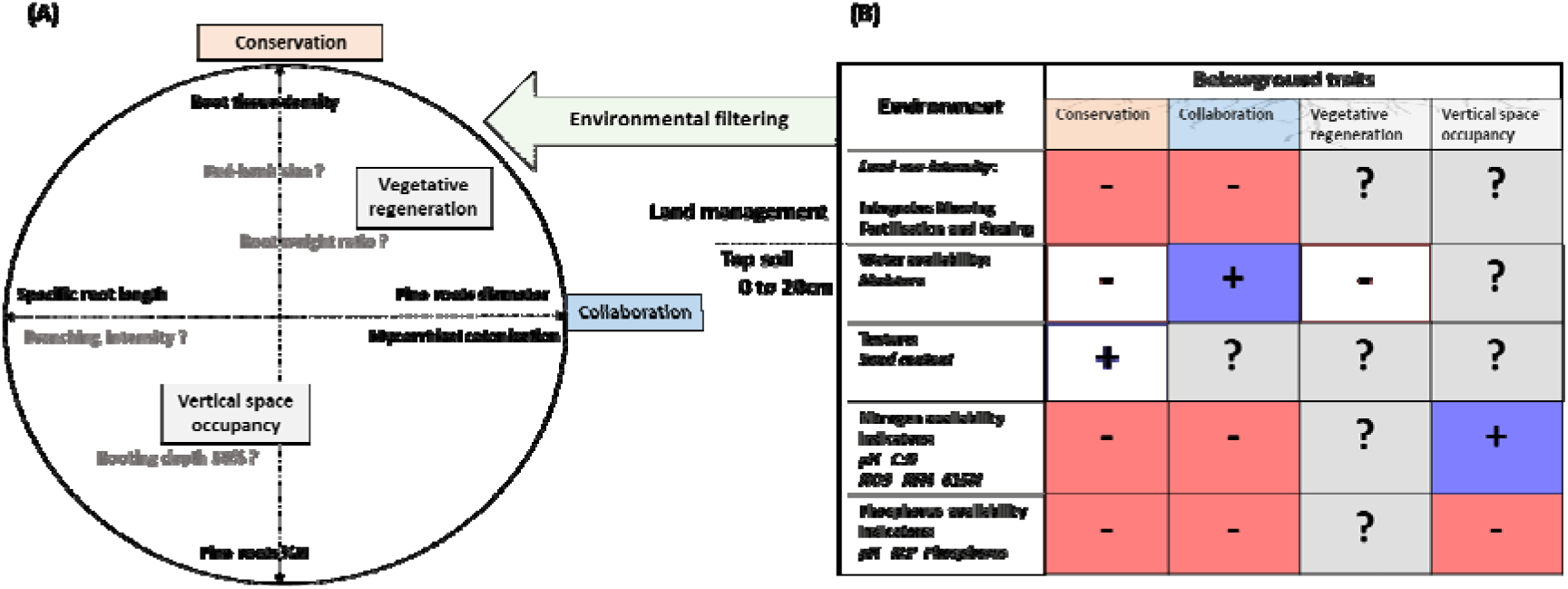
Hypothesized relationships between (A) community weighted means (CWMs) of belowground traits in grasslands potentially aligned on two plant-strategy gradients. A ‘conservation’ and a ‘collaboration’ gradient are expected as the main dimensions of plant variation, as our trait selection contains mostly traits of the root economics space. A ‘vegetative reproduction’ and ‘vertical space occupancy’ aspect could represent additional plant-strategy gradients, or be embedded within the ‘conservation’ and ‘collaboration’ gradients. The two known belowground dimensions are here represented as two orthogonal axes (‘Conservation’ and ‘Collaboration’) with the traits that have previously shown to be associated with them in black font. The positions of four other traits, bud-bank size, root weight ratio, branching intensity and rooting depth 50% (grey font), are yet unknown. (B) As a result of environmental filtering, each plant-strategy gradient could be associated with different or overlapping environmental variables. The signs and colors indicate the hypothesized direction of the relationships. The references on which these hypothesized relationships are based can be found in Appendix S4.

Belowground organs other than roots add another layer of complexity in terms of form and function to plant-trait space. Plant structures such as rhizomes, root buds and tubers play important roles in storage and vegetative reproduction (Klimešová, Martínková, & Ottaviani, 2018). Species with large bud-bank size are more likely to be perennial and ‘slow’ growing (E-Vojtkó et al. 2017). Furthermore, although not strictly a belowground trait, the root-weight ratio, i.e. the proportion of biomass allocated to roots, is a useful indicator of plant investment into the uptake and storage of different resources (Reynolds & D’Antonio, 1996), and may be linked to rooting depth (Schenk and Jackson 2002a) and bud-bank size. It remains to be tested, whether these belowground traits are aligned with the ‘conservation’ or ‘collaboration’ gradient, or rather represent independent plant strategies.

In grassland communities, which cover about 40% of the continents excluding Greenland and Antarctica (Suttie, Reynolds, & Batello, 2005), the few studies on belowground traits and their variation along environmental gradients are generally limited to root morphological traits and only use a limited set of coarse environmental parameters (Craine, Froehle, Tilman, Wedin, & Chapin, 2001; Prieto et al., 2015; Erktan et al., 2018). Analysing the relationships between aboveground traits and various types of environmental factors, such as climate, soil properties and land-use intensity, has already improved our understanding of trait variation in grasslands (Garnier et al., 2007), and should also be applied to belowground traits. In particular, plants may have various strategies to deal with nutrient deficits and imbalances in soils. For example, it is likely that mycorrhizal collaboration becomes more important at soils with limited phosphorus (P) availability (Ma et al., 2020). Similarly the form of plant-available mineral soil nitrogen (ammonium versus nitrate) could also select for different belowground traits, as species vary in their preference for different forms of N (Weigelt, Bol, & Bardgett, 2005; Pornon, Escaravage, & Lamaze, 2007; Maire, Gross, Da Silveira Pontes, Picon-Cochard, & Soussana, 2009).

To better understand how belowground plant traits relate to environmental variation, we investigated 1) how community mean values of different belowground traits align along known plant-strategy gradients, and 2) how the shifts of community means along these strategy gradients depend on environmental variables. Thus, we complemented traits defining the gradients of the root economics space with additional traits that might represent independent strategies of plant functioning. Therefore, we measured traits of plants grown in pot and used database information and vegetation relevés to calculate community weighted means of belowground traits for 150 grassland plots in Germany. We then assessed the dimensionality of the variation in community weighted means of ten traits with principal component analysis (PCA), and related the principal components to ten land-use intensity and soil variables. A priori hypotheses of the relationships between traits, plant strategy dimensions and environmental variables are presented in Fig. 1.

## Methods

### Data on grassland vegetation composition

The plant-community data used as a baseline for Central European mesic grassland vegetation originate from the ‘Biodiversity Exploratories’ project (Fischer et al., 2010). In each of three regions of Germany, the Schwäbische-Alb (south-western Germany), Hainich-Dün (central Germany), and Schorfheide-Chorin (north-eastern Germany), 50 grasslands covering a wide range of land-use intensities were selected. From 2008 to 2019, the vegetation composition of a 4 m × 4 m plot in each of the 150 grasslands was assessed annually in May/June by identifying all vascular plant species and estimating their cover. To align the species names between the vegetation and trait datasets, we standardized the species names according to the accepted names in The Plant List (www.theplantlist.org, accessed 15 June 2019, using the Taxonstand R package (Cayuela, La Granzow-de Cerda, Albuquerque, & Golicher, 2012). In total, 319 vascular plant species have been identified in the 150 grassland plots.

### Plant species traits

We obtained mean species values for eight traits from four pot experiments that we performed, and for two further traits from databases. For 291 of the 319 grassland species, we were able to obtain seeds from commercial seed suppliers or botanical gardens. We then performed four pot experiments to measure species traits. *Taraxacum* spp. are abundant in the grassland plots, though, due to their complex taxonomy, rarely identified at the species level. We here used trait values of *Taraxacum campylodes* for *Taraxacum* spp. The trait values are part of a previously published dataset (Lachaise, Bergmann, Rillig, & van Kleunen, 2020) and an unpublished dataset (Bergmann et al. *unpublished data*), and comprehensive descriptions of the experiments are provided in Appendix S1. In brief, we did one greenhouse experiment in which we grew 2659 individual plants, representing 216 species, for four weeks after which we weighed the roots and analysed scanned images of the roots with WinRHIZO 2017a software (Regent Instruments Inc., Canada) to determine root tissue density, specific root length, fine root diameter, root weight ratio and root branching intensity (Lachaise et al., 2020). We did a second greenhouse experiment using 2007 plants, representing 196 species, to determine the nitrogen content of fine roots (Fine roots %N) using isotope-ratio mass spectrometry. In a third greenhouse pot experiment, we determined mycorrhizal colonization rate for 225 plants, representing 75 species that are among the most common ones in the grasslands plots (mean cover of 65%, Appendix S3). Six weeks after inoculation with spores of *Rhizophagus irregularis* (see Bergmann et al. *unpublished data*), roots were harvested and washed, and the percentage of mycorrhizal colonization was determined using the line-intersect method (McGonigle, Miller, Evans, Fairchild, & Swan, 1990). In a fourth experiment, we grew 752 plants, representing 183 species, in outdoors growth-tubes to determine the depth above and below which plants have 50% of their root biomass (Rooting depth 50%, see Appendix S1 or Schenk & Jackson, 2002 for the calculation method) for about 16 weeks. In addition, to have an estimate of the belowground regeneration potential, we extracted bud-bank size, including stem and root-derived buds occurring belowground or at the soil surface, from the CLO-PLA database (Klimešová, Danihelka, Chrtek, de Bello, & Herben, 2017) for 313 of the 319 species. Finally, to also have a reliable indicator of the plant communities’ acquisitive side of the plant economics spectrum (Allan et al., 2015; Busch et al., 2019), we extracted specific leaf area, the one and only aboveground trait in our analyses, for 279 of the 319 species from the LEDA database (Kleyer et al., 2008).

### Environmental variables of grassland plots

To relate the different dimensions of variation in community weighted trait means of the grassland plots to the abiotic environment, we used ten environmental variables related to land-use intensity and soil conditions. The goal was to capture a relatively independent set of descriptors likely to drive the belowground functioning of plants. A detailed description of each variable can be found in Appendix S2. We used the land-use-intensity index (Blüthgen et al., 2012), which aggregates the intensity of mowing, fertilization and grazing, and is a major driver of ecosystem properties (Allan et al., 2015). We used a variety of physicochemical indicators related to soil fertility of the topsoil (0-20 cm): Soil-moisture content and sand content were measured to capture soil water availability and texture, respectively. Soil pH was chosen, as it affects the availability of essential plant nutrients such as P in soils. We used soil extractable NO_3_, NH_4_ and δ15N as indicators of soil nitrogen availability and related processes (Robinson, 2001; Kleinebecker et al., 2014), and the C:N ratio as a coarse indicator of stoichiometry and organic matter decomposability (Schachtschabep, Blume, Brümmer, Hartge, & Schwertmann, 1998). We further made use of resin-bag-adsorbed phosphorus and the N:P ratio to capture phosphorus availability in soil (Güsewell, 2004). Because soil volume is a central element in soil fertility and root-system distribution, we used data on soil bulk density to convert per-mass nutrient concentrations to per-volume concentrations (Appendix S2). Few of the grassland-site descriptors were measured for each of the years for which we had vegetation-composition data (i.e. for the period 2008-2019). However, we tried to maximize the coverage for this period by using all available census dates for these variables (see Appendix S2 for years covered) and averaging the values per plot.

### Statistical analyses

All the statistics were done using R v 4.0.1 (R Core Team, 2020).

#### Community weighted trait means

To characterize the plant communities of each of the 150 grassland plots based on values of functional traits of their species, we calculated community weighted means (CWMs) as

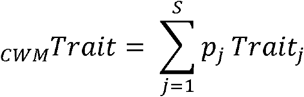

Here *p_j_* is the relative cover of species *j* in the community, Trait*_j_* is the trait value of species *j*, and *S* is the number of species in the community with available trait data. Because some plots had patches of bare soil in some of the annual vegetation surveys, and because for some species trait data were missing, we normalized plant cover to cumulate to 100% for all species with available trait data in each plot before calculating the CWMs. As we have trait data for most of the dominant grassland species, we have data for about 90% of the total cover in most plots, for most traits (Appendix S3). The only exception is mycorrhizal colonization, which was only available for 78 species, but, even for that trait, the average cover of species included was 65% (range 32 - 87%, Appendix S3).

#### Principal components of CWMs variation

As the CWMs of several traits were correlated (Appendix S10), we performed principal component analyses (PCA) to reduce the dimensionality of the data. To assess how robust the resulting dimensions are to the inclusion of additional information, we performed four separate PCAs. Each of these PCAs included all nine belowground traits, but they differed in that we also included or excluded _CWM_*Specific leaf area*, as one of the major traits associated with the aboveground ‘fast’ side of the plant economics spectrum, and that we included or excluded plant-functional-type information (i.e. the percentages cover of graminoids, N-fixing forbs and non-N-fixing forbs). So, one PCA included CWMs of belowground traits only (“Belowground PCA”), one additionally included _CWM_*Specific leaf area* (“Above-Belowground PCA”), one additionally included the proportions of Poales, Fabaceae and non-Fabaceae forbs, and one included all. To increase the separation of the variable loadings (the trait CWMs) on the two first axes, we performed an ‘oblimin’ rotation on these axes for the Belowground PCA and the Above-Belowground PCA. To complement the information provided on taxonomic or phylogenetic influence on community trait values, we also looked at the ten most dominant species or taxa in the trait space formed by PC1 and PC2 and the indicator species or taxa that associated with each quadrant of the two-dimensional space formed by PC1 and PC2 (Appendix S15). As the proportion of plant functional types were shown to be significantly related to specific plant strategies in PC1 and PC2 (Appendix S6, Appendix S10), we also did three additional versions of the Above-Belowground PCA, removing each plant functional type in the CWMs calculation once, to evaluate how much the trait relationships are affected by the presence of the respective plant functional type (Appendix S7). To compare the relationships observed at the community level and at the species level, we also did the Above-Belowground PCA using trait means of the species instead of CWMs (Appendix S16). For each PCA, _CWM_*Root tissue density* was log_10_ transformed and for each trait or proportion of plant functional type, data was standardized by subtracting the mean and dividing by the standard deviation to conform to the multinormality requirements.

#### Associations of the principal components of CWMs with environmental variables

To test for associations between the principal components of CWMs of the grassland plots and the environmental variables, we performed multiple regression. The PC1 and PC2 scores from each of the four PCAs on CWMs of the functional traits were used as response variables, and the environmental variables were used as predictors. C:N, N:P, sand content, NH_4_, NO_3_, and δ15N were log-transformed before analysis to get a more regular (less clumped) distribution of the predictor values. To account for the fact that the grassland plots are located in three different regions of Germany, we also included region as a predictor in the models. For model reduction, backward stepwise model selection based on AIC was performed using the function step(). This procedure selects a parsimonious set of predictors while minimizing the variance inflation factor (max VIF = 3.6 for Above-Belowground PCA). Because the two first axes (PC1 and PC2) of the four PCAs produced similar scores for the CWMs of the grassland plots (all pairwise correlations of the PC1s were >0.98 and those of the PC2s were >0.67), we present the results of the analysis of the “Above-Belowground PCA” in the main text (Fig. 3 based on the PC axes of Fig. 2), and the results for the other three PCAs in Appendix S8. We did the same for the PC3 to PC6 scores from the Above-Belowground PCA (Appendix S13), and for each of the ten _CWM_*Traits* (Appendix S14). We further tested if the proportion of the three plant functional types, as related to the trait dimensions, responded to environmental variables in a similar way, and ran the same models with the proportion of plant functional types as the response variables (Appendix S12).

**Figure 2.**
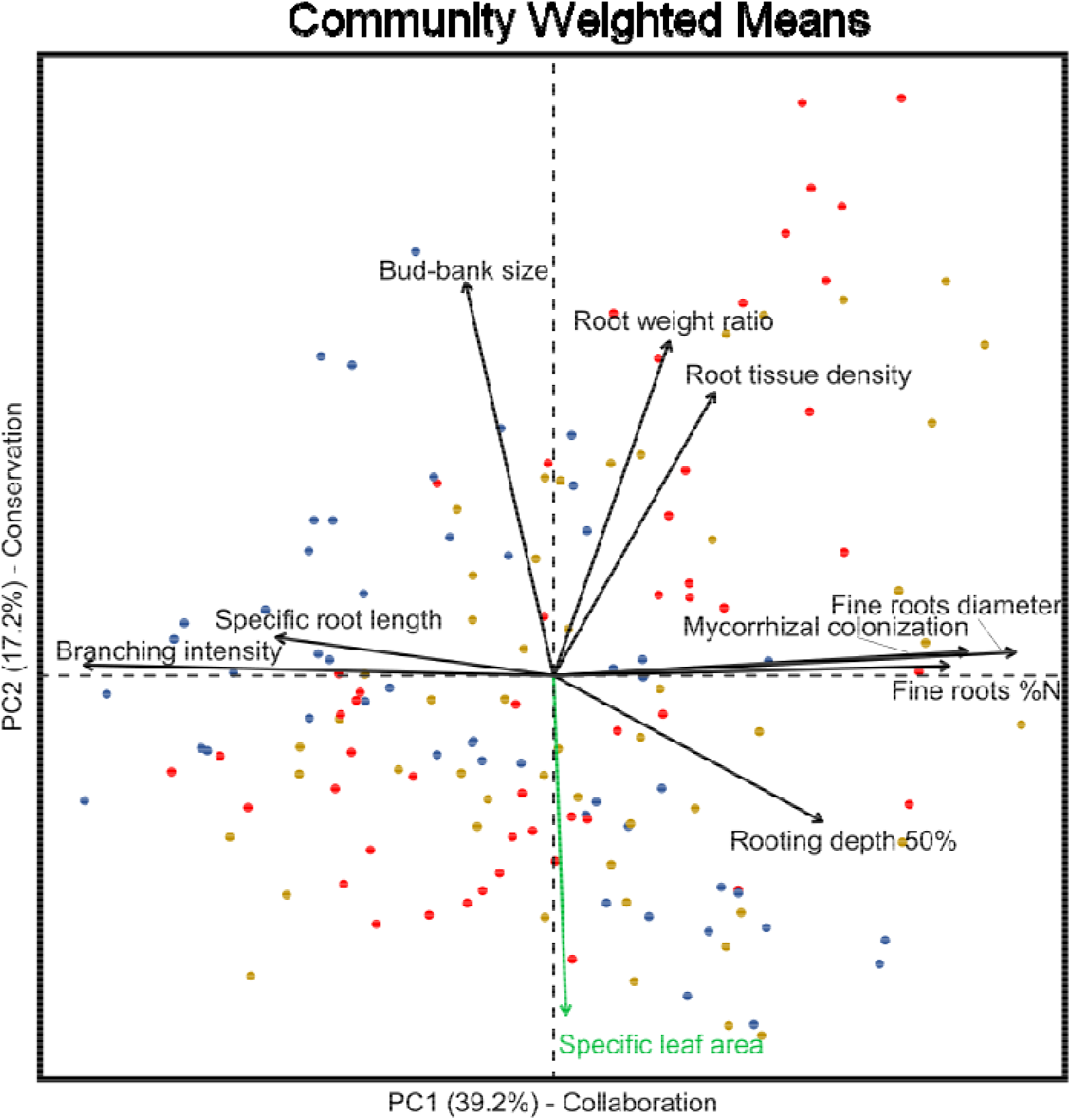
The two first PCs of the Above-belowground PCA, explaining 56.4% of the total variance in community weighted means (CMWs). Every _CWM_*Trait* has a strong loading on either PC (Appendix S9). The subsequent PCs, including PC3 representing 14.8% of the total variance (Appendix S5), mainly captured variation among the three regions, probably capturing differences in regional species pool, but did not strongly relate to any environmental parameter (Appendix S13). The sole aboveground trait that we included, _CWM_*Specific leaf area*, is shown in green. The scores of the 150 grassland plots used for the PCA are shown in different colors for each of the three regions (red for the Schwäbische Alb, brown for Hainich, blue for Schorfheide, each with N=50). PC1 is mostly characterized by CWMs of traits related to the mycorrhizal ‘collaboration’ gradient of the root economic space, with on the left, the ‘do-it-yourself’ strategy and on the right, the ‘outsourcing’ strategy. PC2 is more characterized by CWMs of traits related to the ‘conservation’ gradient of a ‘root and leaf economic spectrum’, with on the top, the ‘slow’ strategy and on the bottom, the ‘fast’ strategy. Bud-bank size, as a surrogate of the vegetative regeneration potential is associated with the ‘slow’ strategy. Correlation coefficients between the CWMs are provided in Appendix S10 and corroborate the relationships observed on PC1 and PC2. The loadings onto PC1 to PC6 (90% of variance explained) are in Appendix S9. To maximize the loadings of the traits characteristic of the ‘collaboration’ and ‘conservation’ gradients on PC1 and PC2, an “oblimin” rotation was performed on the plot scores.

**Figure 3.**
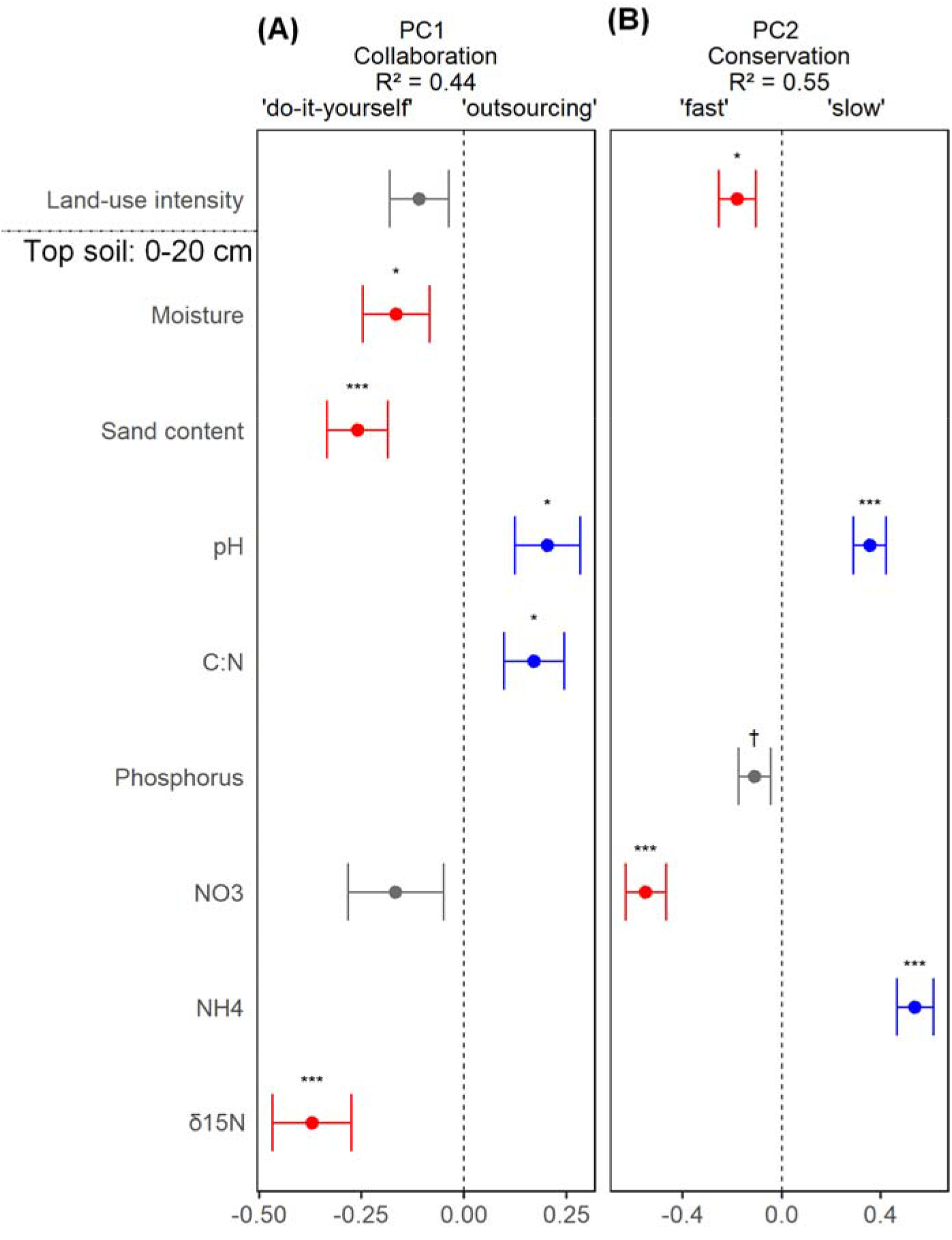
Estimates from linear models testing the effects of environmental variables on PCA scores for (A) PC1 -‘Collaboration’ gradient and (B) PC2 - ‘Conservation’ gradient from the Above-Belowground PCA on community weighted means of traits. On the y-axis are the nine environmental variables that were retained in the most parsimonious models (Region and N:P were not retained). The error bars around the estimates are standard errors. Significant (* for p < 0.05; ** for p < 0.01; *** for p < 0.001) negative and positive estimates are marked in red and blue, respectively. Non-significant (p > 0.05) estimates are marked in grey. Marginally significant (p < 0.10) estimates are marked with †.

## Results

### Dimensionality of CWMs

The Above-Belowground PCA (Fig. 2, Appendix S5) as well as the other three PCAs (Appendix S5, S6) revealed that the two first axes generally explained about 55-60% of the total trait variance, and that each of the 10 traits had intermediate to strong loadings on at least one of these two axes (Appendix S9). PC1 had strong negative loadings of _CWM_*Specific root length* and _CWM_*Branching intensity*, and strong positive loadings of _CWM_*Mycorrhizal colonization*, _CWM_*Fine roots %N*, and _CWM_*Fine roots diameter*. PC2 had strong positive loadings of _CWM_*Bud-bank size*, _CWM_*Root weight ratio*, and _CWM_*Root tissue density*, and strong negative loadings of _CWM_*Specific leaf area*. PC1 thus overall captured the mycorrhizal ‘collaboration’ gradient of the root economic space, and PC2 captured the resource ‘conservation’ gradient. The traits of the two gradients of plant functioning, the ‘collaboration’ and the ‘conservation’ gradients were, however, only partially independent (see CWMs correlations in Appendix S10). The ‘fast’ strategy tended to associate with the ‘do-it-yourself’ strategy. _CWM_*Rooting depth 50%* loaded rather strongly on both of these two PCs (Fig 2A; Appendix S9), suggesting that deep-rooting communities were associated with the ‘outsourcing’ side of the ‘collaboration’ gradient as well as the ‘fast’ side of the ‘conservation’ gradient.

### Associations of the dimensions of CWMs with environmental variables

The position of grassland communities along the ‘collaboration gradient’ (PC1) and the ‘conservation’ gradient (PC2) was significantly related to several environmental variables (Fig. 3). The δ15N isotopic signal, sand content and moisture of the topsoil were associated with the ‘do-it-yourself’ side of the ‘collaboration’ gradient (*i.e*. had negative effects on PC1). Land-use intensity and NO_3_ content were retained by the model-selection procedure, as associating with the ‘do-it-yourself’ side, but their effects were not significant (Fig. 3a). The pH and C:N ratio, on the other hand, were associated with the ‘outsourcing’ side of the ‘collaboration’ gradient (*i.e*. had positive effects on PC1; Fig. 3a).

Among the environmental variables, NO_3_ content and land-use intensity were significantly associated with ‘fast’ communities (*i.e*. had negative effects on PC2; Fig. 3b). Phosphorus content was also associated with ‘fast’ communities, but this effect was only marginally significant (Fig. 3b). NH_4_ content and pH, on the other hand, were significantly associated with the ‘slow’ communities (*i.e*. had positive effects on PC2; Fig. 3b). The effects and the variance explained by the different models are comparable for the four PCAs, with and without _CWM_Specific leaf area and with and without the plant functional types (Appendix S6).

## Discussion

We investigated the belowground trait dimensionality of grassland communities and found that a ‘collaboration’ (do-it-yourself vs. outsourcing) and a ‘conservation’ (slow vs. fast) gradient (*sensu* Bergmann et al. 2020) explained most of the variation in community-weighted means of belowground traits. Three traits that were not considered previously in the belowground trait space were largely part of these two dimensions. _CWM_*Rooting depth 50%* was associated with the ‘outsourcing’ and ‘fast’ strategies, _CWM_*Branching intensity* with the ‘do-it-yourself’ strategy, and _CWM_*Bud-bank size* with the ‘slow’ strategy. _CWM_*Fine root %N* was surprisingly associated with the ‘outsourcing” strategy. Both gradients responded to environmental variables related to soil conditions, and more fertile soils where generally associated with the ‘fast’ and the ‘do-it-yourself’ strategies. In line with this, we also found that land-use intensity was associated with the ‘fast’ strategy and tended to be associated with the ‘do-it-yourself’ strategy.

### Trait relationships and dimensionality of belowground traits

For the grasslands in our study, variation in community weighed means of belowground traits tended to separate along two dimensions that largely correspond to the two ecological root-trait gradients recently identified for species. PC1 related to the collaboration of plants with mycorrhizal fungi. This ‘collaboration’ gradient (Bergmann et al., 2020) ranged from ‘outsourcing’ communities with a high mycorrhizal colonization rate and thick roots but also, surprisingly, high root nitrogen content, to ‘do-it-yourself’ communities with high specific root length and a high root-branching intensity. PC2 related to the construction cost of roots and leaves and the vegetative regeneration potential. This ‘conservation’ gradient ranged from ‘slow’ communities with high root-tissue density, high root-weight ratio and large bud-banks to ‘fast’ communities with high specific leaf area. _CWM_*Rooting depth 50%* relates to both of these PCs, with deep-rooting communities being ‘outsourcing’ and ‘fast’. To maximize the loadings of the traits onto one of the PC axes, we used an ‘oblimin’ rotation, and as a consequence the PCs are not orthogonal to each other. This also shows that the two plant-strategy gradients (i.e. PC1 and PC2) were not entirely independent, as the ‘fast’ strategy and ‘do-it-yourself’ strategy partly coincide, both with PC1 and PC2 scores, and with the traits that are associated with each strategy (Appendix S10). This is partly in accordance with recent findings of Laughlin et al. (2021) who found cold climatic conditions to enhance the probability of occurrence for both ‘fast’ as well as ‘do-it-yourself’ plant species in a global analysis.

At the community level, belowground trait relationships can differ from the one found at the inter-specific level (Craine et al., 2001; Roumet, Urcelay, & Díaz, 2006; Prieto et al., 2015; Schroeder-Georgi et al., 2016; Zhou, Bai, Zhang, & Zhang, 2018; Erktan et al., 2018; Delpiano, Prieto, Loayza, Carvajal, & Squeo, 2020). The trait clustering we found for CWMs was generally in accordance with previous findings of trait clustering among species, both for trees and herbaceous plants (Weemstra et al., 2016; Kramer-Walter et al., 2016; Bergmann et al., 2020). The main exception was _CWM_*Fine roots %N*, which in our study associated with the ‘outsourcing’ side of the collaboration gradient instead of the ‘fast’ side of the conservation gradient. This might be particular to our study using CWMs, as root nitrogen content relates to both the ‘fast’ and the ‘outsourcing’ strategies when we do the PCA at the species level (Appendix S16a), but only with the ‘outsourcing’ strategy when scaling up to community level (Fig. 2). Trait-performance relationships have already been shown to differ between common garden and field condition in the Biodiversity Exploratories grasslands (Breitschwerdt, Jandt, & Bruelheide, 2019). The differences in relationships among species traits and among community weighted means of traits could reflect the multiple constraints exerted by environmental filtering on the trait values selected in a field context. More effort will be required to disentangle filtering effects and phenotypic changes when traits are assessed in controlled versus field conditions.

We found that communities with large bud-banks were on the ‘slow’ side of the ‘conservation’ gradient. Previously, bud-bank size was shown to be rather independent of the plant economics spectrum, as specific leaf area — a key trait in this spectrum — explained less than 2% of variation in bud-bank size among 1359 herbaceous species (Klimešová et al., 2016). In our study, the correlation between bud-bank size and specific leaf area of species mean values was significantly negative (−0.17, p < 0.01; Appendix S16b), though still weaker than between the corresponding CWMs (−0.34; Appendix S10). Because all of our species were selected based on their presence in permanent grasslands, it could be that the association between bud-bank size and ‘conservation’ traits is a feature of this specific habitat. Nevertheless, inclusion of other traits linked to clonality, such as clonal lateral spread, shoot persistence or bud-bank depth could reveal specific clonal strategies (Herben & Klimešová, 2020), potentially increasing the dimensionality of belowground trait space (Ladouceur et al., 2019). The smaller bud-bank size we observed in communities with the ‘fast’ strategy, typical of resource rich grasslands, where competition for light might be more intense (Hautier, Niklaus, & Hector, 2009) could indicate that those plants invest more in immediate aboveground light-harvesting structures at the cost of future regrowth ability. In line with this, we also found that low root weight ratios are indicative of ‘fast’ communities.

### Variation in community-trait dimensions explained by the environment

The ‘collaboration’ and ‘conservation’ gradient in PCA were associated with several environmental variables, partly in an overlapping and partly in a unique manner. About half of the variation in PC1 and PC2 scores was explained by environmental variables. Along the ‘collaboration’ gradient, the ‘outsourcing’ strategy was found on dry, non-acidic soils with a low sand content and low N availability (i.e. high C:N, low δ15N and marginally low NO_3_), and tended to be associated with a low land-use intensity (although not significantly). Along the ‘conservation’ gradient, the ‘slow’ strategy was found on non-acidic soils with low P and NO_3_ but high NH_4_ availabilities, and in sites with low land-use intensities. Hence, although the ‘outsourcing’ and ‘slow’ strategies correspond to two different plant-strategy gradients, they both tend to be associated with relatively unproductive soils under low land-use intensity. In fact, all plots in the related upper right section of the PCA diagram originate from calcareous grasslands on shallow, infertile Rendzic Leptosols that are mostly used as unfertilized sheep pasture and characterized by P or NP-limitation (Klaus et al., 2011).

The relationships we found between the ‘collaboration’ gradient and environmental variables are generally in accordance with the current knowledge in mycorrhizal ecology. For example, (Hempel et al., 2013) found, that among 1752 species of the German flora, obligatory mycorrhizal species tended to be positively associated with dry, non-acidic, infertile habitats. In line with this, we found that drier top soils were associated with deeper rooting, more mycorrhizal-associating communities (Fig. 3, Appendix S14). While mycorrhiza have a well-known positive effect under water limited conditions (Augé, 2001), deeper roots allow the uptake of water from deeper soil layers (Fan, Miguez-Macho, Jobbágy, Jackson, & Otero-Casal, 2017). We found that ‘outsourcing’ communities were also linked with lower δ15N isotopic ratios of the soil. The value of δ15N in the soil is the result of multiple processes implicated in the nitrogen cycle and by the primary source of nitrogen in the system, which can be fixation by legumes and organic or inorganic fertilisation (Robinson, 2001). In previous work at the same plots, a high δ15N has been linked to higher plant productivity and lower species richness, potentially indicating a more open N-cycle with enhanced nitrogen losses (e.g. via leaching) and the dominance of few species with a ‘fast’ strategy (Kleinebecker et al., 2014). Indeed, δ15N is strongly positively correlated with NO_3_ concentration in the soil and moderately with NH_4_, moisture and land-use intensity (Appendix S11). If interpreted as an indicator of more plant-available nitrogen in soils, the negative relationship between δ15N and the ‘outsourcing’ strategy is in line with the finding of reduced mycorrhizal colonization in response to nitrogen addition (Ma et al., 2020) and with our finding that ‘outsourcing’ communities tend to be on the ‘slow’ side of the ‘conservation’ gradient.

As arbuscular mycorrhizal fungi are known to help plants with the uptake of phosphorus, we expected that communities on soils with low phosphorus content would score high on the ‘collaboration’ gradient. Nitrogen addition generally decreases the degree of mycorrhizal colonization in conditions of high P availability and increases it under low P availability at the plot level (Ma et al., 2020). Arbuscular mycorrhizal fungi could also help with nitrogen uptake under conditions of high phosphorus concentrations, with a possible negative relationship between N:P and mycorrhizal colonization rates (Blanke et al., 2005; Blanke et al., 2011). Soils depleted in P or in which P is not plant-available are also selecting root systems with little reliance on mycorrhiza, for example by having cluster roots and carboxylate exudation to mobilize inorganic phosphorus (Lambers, Bishop, Hopper, Laliberté, & Zúñiga-Feest, 2012). Reliance on mycorrhiza could depend on how much phosphorus is available, but also on the balance between phosphorus and nitrogen (i.e. the N:P ratio). Neither the anion-exchange-resin data we used as indicator of soil phosphorus availability nor the N:P ratio was related to the ‘collaboration’ gradient (though N:P was marginally positively associated with _CWM_*Mycorrhizal colonization*, Appendix S14). However, low resin-phosphorus was marginally related to the ‘slow’ side of the ‘conservation’ gradient. The nature of plant-available phosphorus in soil is still debated (Barrow, 2021). Phosphorus is also more available in slightly acidic soils (Alt, Oelmann, Herold, Schrumpf, & Wilcke, 2011). The pH of our soils ranged from acidic to slightly alkaline (min. 4.5, max 7.5), with a mean of 6.5, and only 37 out of 150 plots have a pH between 6.5 and 7 which is often used as an optimum to assess phosphorus availability (Penn & Camberato, 2019). So, the positive effect of pH on the ‘collaboration’ gradient could indicate a lower P availability at high pH values. Furthermore, organic fertiliser, applied in the plots with high land-use intensity, is usually rich in P, and our land-use-intensity variable thus could capture part of the P supply that is not captured by resin-P (r=0.49 between resin-P and land-use intensity, Appendix S11). In conclusion, we did not find a decrease of mycorrhizal colonization when P is more available through the effects of resin-P on ‘collaboration’ gradient (Fig. 3), but we cannot rule out an effect of P availability, because of the potential changes in phosphorus availability through fertilisation and pH changes. The proven P-limitation of plant growth (Klaus et al 2011) in plots allocated in the upper right corner of the PCA-diagram (outsourcing and slow) also points in this direction.

The relationships we found between the ‘conservation’ gradient and environmental variables are overall in line with expectations on how soil fertility should relate to the plant economic spectrum. Accordingly, high land-use-intensity and acidic soils with high phosphorus and nitrate concentrations were associated with the ‘fast’ strategy. The decrease in bud-bank size at higher soil fertility (Fig. 3, Appendix S14) is congruent with recent findings that land-use intensity and nitrogen addition decrease total bud density and rhizome biomass in temperate perennial grasslands (Qian, Wang, Klimešová, Lü, & Zhang, 2021; Ottaviani et al., 2021). Disturbance and habitat-productivity indices of species have been associated with an increase in specific leaf area and a decrease in bud-bank size (Herben, Klimešová, & Chytrý, 2018). In contrast to the negative effect of nitrate concentration on the ‘conservation’ gradient, we found a positive effect of soil ammonium concentration. Species preferences for specific nitrogen forms vary with ecological strategies. Early successional species, which are usually on the ‘fast’ side of the ‘conservation’ gradient, generally prefer nitrate, whereas late successional species, which are usually on the ‘slow’ side, generally prefer ammonium (Britto & Kronzucker, 2002; Warren, 2009). It has also been shown that there might be a trade-off between nitrate and ammonium uptake in grassland species (Maire et al., 2009). Some plants can also inhibit nitrification, thereby retaining NH_4_, which is less prone to leaching than NO_3_ (Boudsocq et al., 2012). High rates of nitrification in fertile soil with high microbial activity could lead to a stronger dominance of nitrogen in form of nitrate. Increase in ammonia oxidation with land-use intensity (and therefore fertilisation) has already been shown in our grassland system (Stempfhuber et al., 2014). So, the positive association between ‘fast’ communities and NO_3_ could reflect an overall higher nitrifying activity of microbial communities in fertile, nitrogen-rich soils. In conclusion, the form of nitrogen available in the soil has contrasting effects on belowground traits, with ammonium being more related to the ‘slow’ strategy and nitrate more related to the ‘fast’ strategy.

### Conclusion

The dimensionality of trait syndromes and their relation to environmental variables are central questions in ecology. At the grassland community level, we found an integration of root branching intensity, root-weight ratio, bud-bank size and rooting depth within the bidimensional ‘collaboration’ and ‘conservation’ trait space previously observed at the species level. The variation of both gradients with environmental variables was partly overlapping and partly independent. Indicators of high soil fertility were generally associated with both the ‘fast’ and the ‘do-it-yourself’ strategies. Overall, our study shows that the belowground plant-strategy gradients identified among species are also applicable to the description of plant communities, and can be linked to environmental variables.

## Supporting information

Appendixes

## Acknowledgments

We thank Otmar Ficht, Maximilian Fuchs and Heinz Vahlenkamp for help setting up the experiments, Beate Rüter, Ekaterina Mamonova, Huy Manh Nguyen, Simon Gommel, Maximilian Rometsch, Anika Schick and Emma Bretherick for help measuring the plant traits. We also thank the managers of the three Biodiversity Exploratories, Konstanz Wells, Swen Renner, Kirsten Reichel-Jung, Sonja Gockel, Kerstin Wiesner, Katrin Lorenzen, Andreas Hemp, Martin Gorke and Miriam Teuscher, and all former managers for their work in maintaining the plot and project infrastructure; Christiane Fischer for giving support through the central office, Andreas Ostrowski for managing the central data base, and Markus Fischer, Eduard Linsenmair, Dominik Hessenmöller, Daniel Prati, Ingo Schöning, François Buscot, Ernst-Detlef Schulze, Wolfgang W. Weisser and the late Elisabeth Kalko for their role in setting up the Biodiversity Exploratories project. The work has been (partly) funded by the DFG Priority Program 1374 “Infrastructure-Biodiversity-Exploratories”. Field work permits were issued by the responsible state environmental offices of Baden-Württemberg, Thüringen, and Brandenburg. We acknowledge funding from the German Research Foundation (DFG, grants KL 1866/12-1 to MvK and 323522591 to MR).

## Declarations

Tom Lachaise performed three of the experiments, ran the analyses and wrote the paper. Joana Bergmann performed one experiment and participated in one of the other three. Matthias Rillig contributed to the design of the experiments. Norbert Hölzel, Valentin Klaus and Till Kleinebecker collected environmental data. Mark van Kleunen designed three of the experiments, advised on data analysis and extensively revised the paper. All authors contributed to revisions. The authors declare no conflicts of interest.

## Data availability

The trait data will be archived in Dryad, and the data DOI will be included at the end of the article. The environmental data is partly publicly available on BExIS and partly under an embargo period of three years. It can be requested directly to the authors of the data (see Appendix S2).

