## Appendixes for "Soil conditions drive belowground trait space in temperate agricultural grasslands"

**Appendix S1** Extended methods for the four experiments on root-trait measurements.

**Experiment on root-system morphology**

From May 1 to October 6, 2017, we performed a glasshouse experiment to measure root-system morphological traits of the study species. As root morphology might depend on nutrient availability, we grew half of the plants per species at an intermediate nutrient level and the other half at a high nutrient level. As plasticity was not of interest for this study, we averaged the trait values per species. Because of the large number of species and the time-consuming measurements, we grew the plants in four temporally shifted (4-6 weeks) batches. We aimed to have each species represented in each batch, and to have a total of seven replicates per species and nutrient level across all batches. Out of the 291 species sown, the seedlings of the species that had germinated (2659 seedlings of 216 species) were transplanted individually into plastic pots (1.3 L) filled with a mixture of sand and vermiculite (1:1 volume ratio). The pots were then randomly allocated to positions in two glasshouse compartments, and allowed to grow for four weeks (night/day 10/14 h; 22/28 ± 1.5 °C; relative humidity 80 ± 15%). Plants were fertilized three times a week with either a low nutrient solution (40 mL with 1500 µM KNO3) or a high nutrient solution (40 mL with 12000 µM KNO3). The fertilizer was a modified version of the Hoagland recipe ([see online](http://tomlachaise.com/wp-content/uploads/2020/12/Modified-Hoagland-solution-for-root-morphology-experiment.pdf)). We grew the plants for four weeks only to avoid roots becoming pot-bound, and to be able to analyze the entire root systems. After carefully washing off the substrate, each root system was cut below the collar and stored separately for <1 week in a plastic tube filled with distilled water at 4°C. Then, root systems were spread individually in a thin layer of water in transparent trays (11 cm × 11 cm) and scanned at 800 dpi with a flatbed scanner modified for root scanning (Epson Expression 10000 XL and 11000 XL). The images were analyzed using the software WinRHIZO^TM^ 2017a (Regent Instruments, Quebec, Canada) to obtain the total root length and root volume. Root systems were then oven-dried for >48 hours at 65 °C and weighed. We calculated specific root length by dividing the total root length by the belowground dry biomass, and root tissue density by dividing the belowground dry biomass by the sum of the root volumes according to Rose, 2017). We estimated branching intensity by dividing the total number of root tips by the total length of the root system. The diameter of fine roots (i.e. distal roots), thought to be the most important roots for nutrient uptake (Freschet & Roumet, 2017) was determined by randomly sampling a distal root branch (or a portion of it) for each root system and calculating the mean of the diameter of the external-internal link obtained with the “Link analysis” function in WinRHIZO. This subsampling excluded the thicker transport roots and allowed to obtain a mean value mostly measured on first order roots. We also dried and weighed the aboveground biomass of each plant, and calculated the root weight ratio (i.e. root biomass divided by total plant biomass).

**Experiment on root nitrogen content (Fine roots %N)**

From 15 January to 8 June 2018, we performed a glasshouse experiment to measure root nitrogen content and ammonium and nitrate uptake rate of the study species. Because of the large number of species and the time-consuming measurements, we grew the plants in three temporally shifted (4–6 weeks) batches. We aimed to have each species represented in each batch, and to have a total of six replicates per species and for each of the three modalities of the N source treatment across all batches. The 2007 seedlings of the species that had germinated (n = 196) were transplanted individually into plastic pots (2 L) filled with a mixture of sand and vermiculite (1:1 volume ratio). The pots were then randomly allocated to positions in two glasshouse compartments, and allowed to grow for six weeks (night/day 10/14 h; 22/28 ± 1.5°C; relative humidity 80 ± 15%). Plants were fertilized three times a week with an intermediate nutrient solution (40 mL with 6000 μM KNO3). The fertilizer was a modified version of the Hoagland recipe ([see online](http://tomlachaise.com/wp-content/uploads/2020/12/Modified-Hoagland-solution-for-nitrogen-uptake-experiment.pdf)). We grew the plants for six weeks to get enough root material for the nitrogen content and uptake analysis. After washing off the substrate between 7.30 and 8.00 am, the root systems were split into three bundles of fine roots. Each bundle was immersed for 120 minutes between 8.00 and 12.00 am in an 8 mL tube containing 4 mL of the following nutrient solutions, which differ only in the form of label used: 1) ^15^N-NO3 labeled 2) ^15^N-NH4 labeled or 3) control solution without ^15^N-label ([see online](http://tomlachaise.com/wp-content/uploads/2020/12/Isotope_experiment_final_solutions_for_incubation.pdf)). After 120 minutes of incubation, immersed fine root bundles were cut off, washed twice with 0.5µM CaCl2, dried with cellulose paper, and stored at −20°C until their fresh weight determination. They were then oven-dried for >48 h at 65°C, weighed, and finely ground. Fine root materials (0.5–1.5mg aliquots) were processed in an Elemental Analyser (Elementar, Analysensysteme, Germany) to determine total nitrogen content (Fine roots %N).

**Experiment on mycorrhizal colonization rate**

All data on mycorrhizal colonization originate from a pot experiment that took place in a controlled greenhouse in Berlin in 2018 (16 h daylight at 22°C, 8 h night at 15 °C). The study included a larger species set but only 75 of them occurred in the grassland 8 replicates were set up per species distributed over 4 time blocks. Dead replicates were substituted it in the following time block if necessary. Seeds were surface sterilized once before the experiment (3 min in 7% bleach, washing in de-ionized (DI) water), dried at 20 °C and stored for 2-12 weeks until sowing. Germination took place in plastic boxes on 1:1 steamed sand and vermiculite (1-3 mm, ISOLA Vermiculite GmbH; Sprockhövel, Germany). Seeds were sown consecutively based on pre-findings to assure that seedlings were between the cotyledon stage the stage of first leaves at time of transplanting. The actual experiment took place in plastic cones (410 mL) filled with the same substrate as for germination.

Cones first got filled to c. ¾. Subsequently we added a 30 mL horizon of a 1:1 mixture of steamed sand and mycorrhizal inoculum in 1-2 mm vermiculite (INOQ Agri, Inoq GmbH, Schnega, Germany). According to supplier information, the inoculum contains 145 spores/mL of *Rhizophagus irregularis*, a mycorrhizal fungus commonly used in agricultural and scientific approaches in temperate habitats (Lenoir et al. 2016). The horizons were further covered with ~30 mL substrate. Seedlings received 30 mL of DI water during transplantation. We allowed dead seedlings to be replaced during the first week. Plants grew for 6 weeks within each time block. At time of transplanting all pots were fully randomized and got rearranged every two weeks. Plants received 25 mL of DI water 3 times a week, plants were watered with 25 mL of DI water while instead 25 mL of a ¼ strength Hoagland solution was applied two weeks and four weeks after transplanting. At time of harvest, roots were carefully washed by hand and kept in water at 4°C for less than a week before they got dried at 60°C for three days. For the determination of the mycorrhizal colonization, we used representative subsamples of the total root system of three random replicates per species. Dry roots were cut into small pieces and cleared in 10% KOH for 15 min at 80°C followed by staining in 0.05% Trypan Blue in Lactoglycerol for another 15 min at 80°C. We determined mycorrhizal colonization with the 200× magnified intersection method (McGonigle *et al.*, 1990). Each slide contained a minimum of 30 root pieces to count presence or absence of mycorrhizal structures in 50-100 intersects. Mycorrhizal colonization rates of up to 86% confirmed a successful inoculation.

**Experiment on rooting depth**

From the 15th of May to the 10th of October 2018, we performed an outdoor pot experiment to measure the maximum rooting depth of the species. Up to five seedlings of the species that had germinated (N=183 out of the 291 species sown), totaling 752 plants, were transplanted individually into 120 cm high plastic tree shelter tubes (Tubex ® Standard Plus, http://www.tubex.com/products/tree-shelters/tubex-standard-treeshelters/specification.php), which are normally used in forestry to protect young trees against animals and the elements. We closed the bottoms of these tubes with thick pieces of cotton tissue to be able to use them as pots. The tubes were filled with a mixture of sand and vermiculite (1:1 volume ratio) up to a height of 115 cm. This substrate can be easily penetrated by the roots, and therefore allows each species to reach its maximum rooting depth quickly. The tree shelter tubes were delivered in packages of five tubes stacked into one another, and they, therefore, came in five diameter classes (8.0, 8.4, 10.0, 10.8, and 12.0 cm). To avoid that tube diameter would be confounded with species identity, each of the five seedlings per species was planted in a different tube-diameter class. We placed the tubes upright in a randomized design in the Botanical Garden of the University of Konstanz (47° 41' 24.0" N 9° 10' 48.0").

We planted 752 plants but, due to early mortality, we had to replace 126 of them within the next three weeks. The growth period, therefore, ranged from 16 to 19 weeks. The experiment took place during the summer of 2018 (mean temperature: 19.5 °C, min/max 2.5/37.4 °C; relative humidity: mean 74%, min/max 22.7/100%). All the plants were fertilized once a week with 60 mL of a standard nutrient solution (1‰ Universol® Blue, Nordhorn, Germany), and watered regularly from above. We harvested the plants in October 2018. Each tube was carefully sliced open, and we measured the distance from the top of the substrate to the deepest root. During the harvest of each plant root system, four slices of different depths (0-15 cm; 15-30 cm; 30-60 cm; 60-115 cm) were performed, and the root biomass contained in each slice was separately weighted. To adjust for individual plant maximum rooting depth, the depth of the last slice was stopped at the deepest observed root. For example, if the maximum rooting depth of a plant was 50 cm, then the corresponding last slice of the plant was assigned to the depth of 30-50 cm. Then for each plant, a model of exponential decrease of biomass with depth was fitted (Schenk & Jackson, 2002). As the whole root system was sampled, we could interpolate the depth at which 50% of the total root system biomass was located. Compared to the maximum rooting depth, which is the distance from the top of the substrate to the deepest root, the rooting depth 50% takes into account the difference in biomass investment at different depths. This should better capture the depth at which most water and nutrient uptake take place. The two metrics are however strongly correlated (Pearson’s r = 0.78).

**Appendix S2** Information about the environmental variables of the Biodiversity Exploratories grassland plots used in the linear regression models presented in Fig. 3, Appendix S8, S12, S13, and S14. The data were collected in the period 2008-2019, corresponding to the period covered by the CWMs calculation from vegetation surveys. The datasets are present in the [BExIS database](https://www.bexis.uni-jena.de/) https://www.bexis.uni-jena.de/ (http://doi.org/10.17616/R32P9Q). Some of the datasets are already public and some are still under the three years embargo period. A description of the metadata provided by their authors of each dataset is available below. It was visually assessed that most of the variation in soil variables for which several years of data are available comes from differences among plots rather than differences over time. Soil depth and bulk density were not used directly in our models and are indicated in grey. Soil-depth measurements were performed in only 144 plots and were constrained to a maximum depth of 100 cm, leading to an underestimation of its variance and a considerable bias in the estimation of its true value in the Schorfheide region, where soils are generally deeper than in the other two study regions. Bulk density was strongly correlated to many other soil variables and was used to convert the soil variables originally expressed per unit of soil mass to the following units of volume: sand content and phosphorus in g/m3 and mg/m3 of soil respectively, C:N, N:P, NH_4_, NO_3_, δ15N per cm^3^ of soil instead of per g of soil.

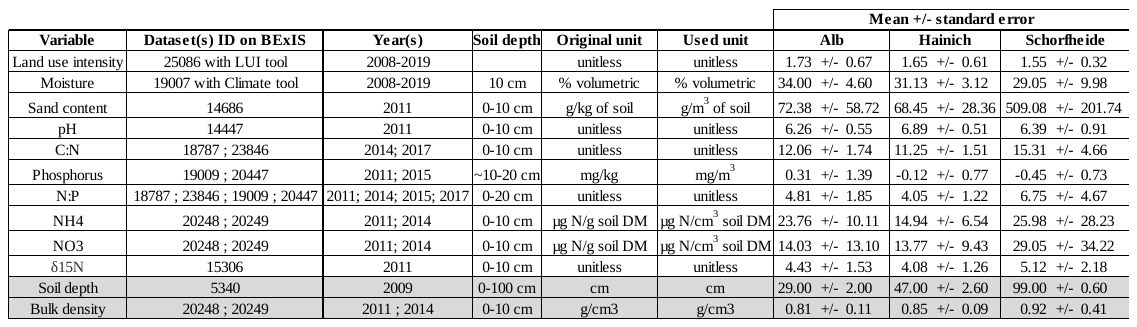

| **Dataset ID in BExIS** | **Used for variable(s)** | **Description provided by data authors in BExIS metadata and published studies as of 25 February 2021 (edited for readability)** |
| --- | --- | --- |
| 25086 | Land-use intensity | The data derive from values of the land-use survey (annual interviews with all grassland land owners / users of the three Exploratories basic Dataset 23746). These data are the land management components (mowing, grazing, fertilization) for the LUI Calculation Tool.  **Calibration**: Livestock unit conversion factor: cattle: <0.5 years = 0.3, 0.5-2y =0.6, >2y = 1 sheep/goats: <1y = 0.05, >1y = 0.1 ponies/small horses: 0.7 horses: <3y = 0.7, > 3y = 1.1 Conversion factor for total nitrogen input: Manure [t/ha] with conversion factor [kg/t]: cattle: 5.6, horse: 4.9, sheep: 8.13 Slurry [m³/ha] with conversion factor [kg/m³]: cattle: 3.85, pig 5.4, mixed: 4.45, biogas/digestate: 4.  **Procedure**: Formulae calculated per year and plot: mowing = number of cuts grazing = livestock units*grazing days/ha summed up for all grazing periods livestock units consist of the number of livestock multiplied by a conversion factor, which is species and age dependent (see above). Fertilization = amount of applied nitrogen within the fertilizer per ha. The nitrogen content depends on the type of fertilizer.  ***Main reference(s):*** Blüthgen, N., Dormann, C. F., Prati, D., Klaus, V. H., Kleinebecker, T., Hölzel, N., . . . Weisser, W. W. (2012). A quantitative index of land-use intensity in grasslands: Integrating mowing, grazing and fertilization. Basic and Applied Ecology, 13(3), 207–220. https://doi.org/10.1016/j.baae.2012.04.001 |
| 19007 | Moisture | Climate Data Set - Time Series Web Interface  Automated environmental monitoring by including air pressure, wind, precipitation and radiation at 10 min resolution - aggregated to 1hour + 1day + 1week + 1month + 1year.  **Instruments**: ADL-MX Datalogger System; DeltaT ML2X Soil Humidity Probe; MNT FExtension 2010 Soil and Ground Surface Temperature Sensor; MELA KPC1/5-ME & #IAK1.00.F137.520.CS8  Procedure: quality 0 - no quality check  quality 1 - just physical range check  quality 2 - physical range + step range check  quality 3 - physical range + step + empirical check  ***Authors Procedure:*** Hourly aggregation was used for soil moisture for the time series 2008-2019 |
| 14686 | Sand content | Soil texture focuses on the soil particles that are less than 2 mm in diameter. It indicates the relative content of sand (2-0.063 mm), silt (0.063-0.002 mm) and clay (<0.002 mm). Size separation of particles is based on sieving (particles >0.063 mm) and on the principle that a solution settles out at a rate that depends on the size of the particles. The larger the particle size, the faster particles settle. The settling rate is given by Stokes Law. Soil texture tends to be relatively stable over time since weathering only very slowly changes particle size distribution in soil.  Texture analysis consists of three main steps: (i) destruction of soil organic matter with hydrogen peroxide, (ii) dispersion of soil aggregates into discrete units, and (iii) separation of soil particles of different size by sieving and sedimentation (DIN-ISO 11277).  **Instruments:** 0.63, 0,2, 0.063 mm sieves for isolation of sand fractions (Retsch, Haan, Germany). Instrumentation for pipette method including Atterberg cylinders to separate silt and clay fractions  **Calibration:** Reference soil of VDLUFA/LUFA for quality control (LUFA, Speyer, Germany)  **Procedure:** In each of the 300 experimental plots of the biodiversity exploratories we collected 14 soil cores with a split tube sampler (diameter of 5 cm) along two 20 m transects in grasslands and 40 m transects in forests in May 2011. Organic layers in forests and aboveground plant parts in grasslands were removed before coring. We then prepared a composite sample from the 14 soil cores by mixing the upper 10 cm of the soil. A subsample of the composite sample was sieved to <2 mm, dried and subsequently used for soil texture analysis.  ***Main reference(s):*** Birkhofer, K., Schöning, I., Alt, F., Herold, N., Klarner, B., Maraun, M., . . . Schrumpf, M. (2012). General relationships between abiotic soil properties and soil biota across spatial scales and different land-use types. PloS One, 7(8), e43292. https://doi.org/10.1371/journal.pone.0043292 |
| 14447 | pH | The soil pH was measured in a weak (0.01 M) calcium chloride solution using a pH meter.  **Instruments:** WTW pH meter 538 (WTW, Weilheim, Germany)  WTW pH glass electrode SenTix 61 (WTW, Weilheim, Germany)  **Calibration:** The pH meter was calibrated using buffer solutions with a pH of 4 and 7.  **Procedure:** In each of the 300 experimental plots of the biodiversity exploratories we collected 14 soil cores with a split tube sampler (diameter of 5 cm) along two 20 m transects in grasslands and 40 m transects in forests in May 2011. Organic layers in forests and aboveground plant parts in grasslands were removed before coring. We then prepared a composite sample from the 14 soil cores by mixing the upper 10 cm of the soil. Soil samples were sieved to <2 mm and air-dried. Subsequently, 10 g of sieved and air-dried soil were mixed with 25 mL 0.01 M CaCl2 solution and shook for 2 hours. Afterwards the pH of the soil suspension was measured using a glass electrode. The pH of each sample was measured twice (pH 1 and pH 2).  ***Main reference(s):*** Birkhofer, K., Schöning, I., Alt, F., Herold, N., Klarner, B., Maraun, M., . . . Schrumpf, M. (2012). General relationships between abiotic soil properties and soil biota across spatial scales and different land-use types. PloS One, 7(8), e43292. https://doi.org/10.1371/journal.pone.0043292 |
| 18787 | C:N ; N:P | Nitrogen occurs as organic and inorganic N in soils. With the CN analysis, however, only the total N is determined. The C:N ratio in soil is calculated as the mass of organic carbon to the mass of total nitrogen. The C:N ratio is as well-established indicator for the nitrogen availability in soil. Total C (TC) and total N (TN) concentrations in soil were determined using an elemental analyser. After removal of organic carbon (OC) by ignition of soil samples at 450 °C for 16 h, inorganic C (IC) was determined with the same elemental analyzer. OC concentrations were calculated as the difference between TC and IC.  **Instruments:** Elemental Analyser (VarioMax, Elementar, Hanau, Germany)  **Calibration:** The C and N analyser was calibrated with glutamic acid.  **Procedure:** In each of the 300 experimental plots of the biodiversity exploratories we collected 14 soil cores with a split tube sampler (diameter of 5 cm) along two 20 m transects in grasslands and 40 m transects in forests in May 2014. Organic layers in forests and aboveground plant parts in grasslands were removed before coring. We then prepared a composite sample from the 14 soil cores by mixing the upper 10 cm of the soil. Soil samples were sieved to <2 mm and air-dried. Subsequently, subsamples of sieved soil were ground in a ball mill. Total C and N concentrations were determined by dry combustion in an elemental analyser at a temperature of 1100°C. The evolving CO2 and N2 was determined with a Thermal Conductivity Detector (TCD).  ***Authors procedure:*** For soil N:P calculation, total soil N was averaged across sampling in 2014 (dataset 18787) and 2017 (dataset 23846) and was divided by Resin P collected in 2011 (dataset 19009) and 2015 (dataset 20447)  ***Main reference(s):*** Birkhofer, K., Schöning, I., Alt, F., Herold, N., Klarner, B., Maraun, M., . . . Schrumpf, M. (2012). General relationships between abiotic soil properties and soil biota across spatial scales and different land-use types. PloS One, 7(8), e43292. https://doi.org/10.1371/journal.pone.0043292 |
| 23846 | C:N ; N:P | The C:N ratio in soil is calculated as the mass of organic carbon to the mass of total nitrogen. The C:N ratio is as well-established indicator for the nitrogen availability in soil. Total C (TC) and total N (TN) concentrations in soil were determined using an elemental analyser. After removal of organic carbon (OC) by ignition of soil samples at 450 °C for 16 h, inorganic C (IC) was determined with the same elemental analyzer. OC concentrations were calculated as the difference between TC and IC.  **Instruments:** Elemental Analyser (VarioMax, Elementar, Hanau, Germany)  **Calibration:** The C and N analyser was calibrated with glutamic acid.  **Procedure:** In each of the 300 experimental plots of the biodiversity exploratories we collected 14 soil cores with a split tube sampler (diameter of 5 cm) along two 20 m transects in grasslands and 40 m transects in forests in May 2017. Organic layers in forests and aboveground plant parts in grasslands were removed before coring. We then prepared a composite sample from the 14 soil cores by mixing the upper 10 cm of the soil. Soil samples were sieved to <2 mm and air-dried. Subsequently, subsamples of sieved soil were ground in a ball mill. Total C and N concentrations were determined by dry combustion in an elemental analyser at a temperature of 1100°C. The evolving CO2 and N2 was determined with a Thermal Conductivity Detector (TCD).  ***Authors procedure:*** For soil N:P calculation, total soil N was averaged across sampling in 2014 (dataset 18787) and 2017 (dataset 23846) and was divided by Resin P collected in 2011 (dataset 19009) and 2015 (dataset 20447)  ***Main reference(s):*** Birkhofer, K., Schöning, I., Alt, F., Herold, N., Klarner, B., Maraun, M., . . . Schrumpf, M. (2012). General relationships between abiotic soil properties and soil biota across spatial scales and different land-use types. PloS One, 7(8), e43292. https://doi.org/10.1371/journal.pone.0043292 |
| 19009 | Phosphorus ; N:P | **Procedure:** Different P fractions of 0.5 g air-dried soil samples were measured after a method described by Hedley et al. (1982) modified by Kuo (1996). The first fraction extracted with anion exchange resins (‘‘resin-P’’) was not analyzed separately but is included in the NaHCO3 extractable fraction. The sequential extraction scheme had four steps (NaHCO3-P, NaOH-P, HCl-P, H2SO4-P). First, 20mL 0.5 M NaHCO3 (pH= 8.5) were added to the sample and the suspension was shaken for 30 min, decanted, and filtrated. Second, 30mL 0.1 M NaOH were added to the remaining soil followed by shaking for 16 h, decantation and filtration. Third, the remaining soil was mixed with 0.1MHCl, heated in a water bath (80°C, 30 min), and cooled down (1 h), followed again by decantation and filtration. Fourth, the residual soil mass was stored overnight in a porcelain crucible in a muffle furnace at 550°C to destroy all organic material. Thereafter, 20mL 0.5MH2SO4 were added and the suspension was shaken for 16 h, decanted, and filtrated. The decanted solution of each step was analyzed for P concentrations. All Pi contents were analyzed using the ammonium molybdate-ascorbic acid blue method described by Murphy and Riley (1962) and measured with a continuous flow analyzer (CFA, AA3, XY2, Seal-Analytic, Norderstedt, Germany) at l=660 nm. Total dissolved P in NaHCO3-extract and NaOH extract was measured with Inductively Coupled Plasma/Optical Emission Spectrometry (ICP-OES, PerkinElmer Optima 5300 DV, S10 auto sampler, l=213.617 nm). For the labile and moderately fractions (NaHCO3-P, NaOH-P), Po was calculated by subtracting Pi from total dissolved P concentrations in the extracts. In addition to the individual fractions, inorganic P in the labile and moderately labile fraction was calculated as the sum of NaHCO3-Pi and NaOH-Pi (SNaHCO3-Pi + NaOH-Pi) and organic P in the labile and moderately labile fraction as the sum of NaHCO3-Po and NaOH-Po (SNaHCO3-Po + NaOH-Po). Total P concentrations represent the sum of all P fractions.  **Instruments:** Continuous Flow Analyzer (SEAL Analytical, Norderstedt, Germany)  Murphy J & Riley JP (1962): A modified single solution method for the determination of phosphate in natural waters. Anal. Chim. Acta, 27, 31-36.  **Main references:**  McLaughlin MJ, Alston AM & Martin JK (1986): Measurement of phosphorus in the soil microbial biomass: A modified procedure for field soils. Soil Biol. Biochem., 18, 437-443.  Kuo S (1996): Phosphorus. In: eds. Sparks DL, Page AL, Helmke PA, Loeppert R, Soltanpour PN, Tabatabai MA, Johnston AE, Sumner ME, Methods of Soil Analysis. Part 3 - Chemical Methods, 5, 869-919 pp. Madison, Wisconsin: SSSA.  Sorkau, Elisabeth; Boch, Steffen; Boeddinghaus, Runa S.; Bonkowski, Michael; Fischer, Markus; Kandeler, Ellen et al. (2018): The role of soil chemical properties, land use and plant diversity for microbial phosphorus in forest and grassland soils. In *J. Plant Nutr. Soil Sci.* 181 (2), pp. 185–197. DOI: 10.1002/jpln.201700082.  ***Authors procedure:*** For soil N:P calculation, total soil N was averaged across sampling in 2014 (dataset 18787) and 2017 (dataset 23846) and was divided by Resin P collected in 2011 (dataset 19009) and 2015 (dataset 20447) |
| 20447 | Phosphorus ; N:P | **Procedure:** Soil nutrient concentrations were measured in situ using 1326 IER bags, made out of nylon fabric containing anion/cation mixed-bed resin beads plus specific resin beads for anionic heavy metals and phosphate (TerrAquat GmbH, Nürtingen; Mischbettaustauscher). Bags contained in total 19.5 g (dry weight) resin and were of a round shape with a diameter of 5 cm. Three bags per plot/subplot were installed in all grasslands for approximately 145 days from March to early August 2015. Three bags were buried at each grassland in 20 cm depth. In the study regions Schwäbische Alb and Hainich, at some very shallow soils, bags could be installed in 10–15 cm depth only.  After removal, IER bags were stored in a fridge at 4°C. Extraction was done for each bag separately. About 15 g of resin was extracted with 100 ml 1 M NaCl, in two steps of two parallels of 50 ml each, shaken for 30 min and filtered. The measurement of NH_4_–N and NO_3_–N was performed with a Continuous Flow Auto Analyser (Skalar Analytic B.V., Breda, the Netherlands). To determine PO_4_–P another aliquot of the resin beads (15 g) was extracted with 100 ml 0.5 M H_2_SO_4_ following the same protocol as described above. All concentrations are given as mean values per plot in mg/g (dry weight) resin. One observational and two experimentally treated grasslands had to be excluded from the study for technical reasons. See Klaus et al. (2018) for further details.  ***Main reference(s):***  Klaus, V. H., Kleinebecker, T., Busch, V., Fischer, M., Hölzel, N., Nowak, S., ... & Hamer, U. (2018). Land use intensity, rather than plant species richness, affects the leaching risk of multiple nutrients from permanent grasslands. Global Change Biology, 24(7), 2828-2840.  ***Authors procedure:*** For soil N:P calculation, total soil N was averaged across sampling in 2014 (dataset 18787) and 2017 (dataset 23846) and was divided by Resin P collected in 2011 (dataset 19009) and 2015 (dataset 20447) |
| 20248 | NH4 ; NO3 | Within the joint soil sampling campaign 2011 all samples are taken in the beginning of May 2011 as a mixed sample from 14 soil cores of the top horizon (0-10 cm) of the all 50 grassland experimental plots of all three exploratories (AEG1–AEG50; HEG1-HEG50; SEG1-SEG50).  **Procedure:** Based on the non-fumigated samples of the Chloroform-fumigation-extraction method (CFE), according to Vance et al. (1987) and Keil et al. (2011), the extractable organic carbon (EOC) and extractable nitrogen (EN) were determined. In brief, C and N were extracted from each non-fumigated replicate (5 g) with 40 mL 0.5 M K2SO4. The suspension was horizontally shaken (30 Min, 150 rpm) and centrifuged (30 Min, 4400 x g). C and N concentrations in dissolved (1:4, extract:deion. H2O) extracts were measured with a TOC/TN analyzer (Multi N/C 2100S, Analytik Jena AG, Jena, Germany). Mineral nitrogen in form of ammonium (NH4+) and nitrate (NO3-) was determined by DIN ISO 14256-2 (2006) with an AutoAnalyzer 3 (Bran & Luebbe, Norderstedt, Germany) on the undiluted extracts from CFE analysis.  ***Main reference(s):***  Keil, Daniel; Meyer, Annabel; Berner, Doreen; Poll, Christian; Schützenmeister, André; Piepho, Hans-Peter et al. (2011): Influence of land-use intensity on the spatial distribution of N-cycling microorganisms in grassland soils. In *FEMS Microbiol Ecol* 77 (1), pp. 95–106. DOI: 10.1111/j.1574-6941.2011.01091.x.  Vance, E. D.; Brookes, P. C.; Jenkinson, D. S. (1987): An extraction method for measuring soil microbial biomass C. In Soil Biology and Biochemistry 19 (6), pp. 703–707. DOI: 10.1016/0038-0717(87)90052-6. |
| 20249 | NH4 ; NO3 | Within the joint soil sampling campaign 2014 all samples are taken in the beginning of May 2014 as a mixed sample from 14 soil cores of the top horizon (0-10 cm) of the all 50 grassland experimental plots of all three exploratories (AEG1–AEG50; HEG1-HEG50; SEG1-SEG50).  **Procedure:** Based on the non-fumigated samples of the Chloroform-fumigation-extraction method (CFE), according to Vance et al. (1987) and Keil et al. (2011), the extractable organic carbon (EOC) and extractable nitrogen (EN) were determined. In brief, C and N were extracted from each non-fumigated replicate (5 g) with 40 mL 0.5 M K2SO4. The suspension was horizontally shaken (30 Min, 150 rpm) and centrifuged (30 Min, 4400 x g). C and N concentrations in dissolved (1:4, extract:deion. H2O) extracts were measured with a TOC/TN analyzer (Multi N/C 2100S, Analytik Jena AG, Jena, Germany). Mineral nitrogen in form of ammonium (NH4+) and nitrate (NO3-) was determined by DIN ISO 14256-2 (2006) with an AutoAnalyzer 3 (Bran & Luebbe, Norderstedt, Germany) on the undiluted extracts from CFE analysis.  ***Main reference(s):***  Keil, Daniel; Meyer, Annabel; Berner, Doreen; Poll, Christian; Schützenmeister, André; Piepho, Hans-Peter et al. (2011): Influence of land-use intensity on the spatial distribution of N-cycling microorganisms in grassland soils. In *FEMS Microbiol Ecol* 77 (1), pp. 95–106. DOI: 10.1111/j.1574-6941.2011.01091.x.  Vance, E. D.; Brookes, P. C.; Jenkinson, D. S. (1987): An extraction method for measuring soil microbial biomass C. In Soil Biology and Biochemistry 19 (6), pp. 703–707. DOI: 10.1016/0038-0717(87)90052-6. |
| 15306 | δ^15^N of soil samples | **Procedure:** Soil sampling was conducted in early May 2011. On each plot, composite samples of 14 topsoil cores (0–10 cm depth) were collected using a split-tube sampler with a diameter of 5 cm. Cores were taken along two 20-m transects across the respective plot, and roots were removed from the samples in the field. Soil samples were air-dried, sieved to < 2 mm and ground to fine powder. An aliquot of 2 mg of the ground samples was then analysed for stable isotope ratio (δ^15^N) using a continuous flow stable isotope ratio mass spectrometer (Finnigan MAT DeltaPlus attached to a Carlo Erba Elementar Analysator with ConFlo II Interface).  ***Main reference(s):***  Kleinebecker, T., Hölzel, N., Prati, D., Schmitt, B., Fischer, M., & Klaus, V. H. (2014). Evidence from the real world: 15 N natural abundances reveal enhanced nitrogen use at high plant diversity in Central European grasslands. Journal of Ecology, 102(2), 456–465. https://doi.org/10.1111/1365-2745.12202 |

**Appendix S3** Number of species used for the calculation of community weighted means (CWMs) for traits and proportions for plant groups, and mean proportions of cover for CWMs and plant groups across the 150 grassland plots.

| **Community parameter** | **Number of species** | **Mean cover** |
| --- | --- | --- |
| _CWM_*Specific leaf area* | 279 | 95% |
| _CWM_*Root weight ratio* | 216 | 92% |
| _CWM_*Bud-bank size* | 313 | 91% |
| _CWM_*Root tissue density* | 216 | 92% |
| _CWM_*Specific root length* | 216 | 92% |
| _CWM_*Root branching intensity* | 216 | 92% |
| _CWM_*Fine roots diameter* | 216 | 92% |
| _CWM_*Mycorrhizal colonization* | 75 | 65% |
| _CWM_*Fine roots %N* | 196 | 89% |
| _CWM_*Rooting depth 50%* | 183 | 90% |
| Proportion of Poales | 40 | 57% |
| Proportion of Forbs | 254 | 30% |
| Proportion of Fabaceae | 31 | 9% |
| Proportion of Annuals | 80 | 3% |
| Proportion of Multiannuals | 246 | 95% |

**Appendix S4** References for the expected relationships between potential dimensions of belowground economic space and environmental variables. We combined results from meta-analysis, field and greenhouse experiments, and plant nutrition manuals to formulate hypotheses on the existence and the direction of a relationship between a belowground functional aspect and an environmental factor. The direction of the relationship is indicated with a “+” when positive, with a “-” when negative, and with “+/-” for mixed expectations regarding the direction. Directional relationships with a strong degree of confidence are shown in red, with low degree of confidence in orange, and with mixed results regarding the direction in grey. The studies included focussed overwhelmingly though not exclusively on herbaceous communities. This table does not intend to provide a comprehensive review of the relationships between belowground traits and environment, but rather to emphasize that different plant-strategy gradients or aspects could be influenced by the same or different environmental drivers.

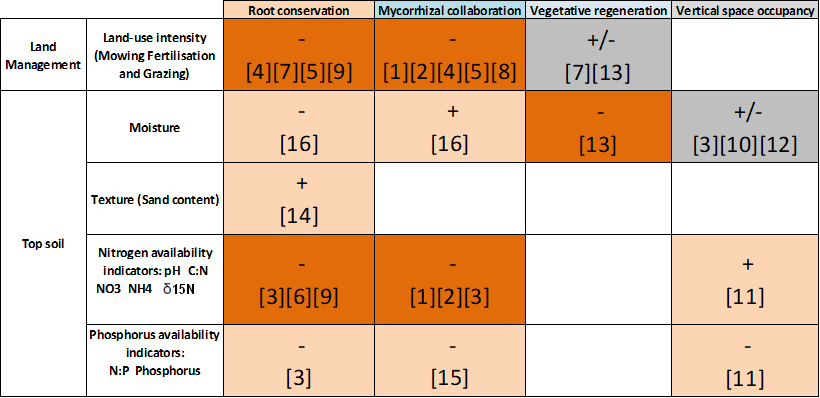

|  | **References for Appendix S4** |
| --- | --- |
|  | [1] Han, Y. et al. 2020. Responses of arbuscular mycorrhizal fungi to nitrogen addition: a meta‐analysis. - Glob Change Biol. |
|  | [2] Ma, X. et al. 2020. Global negative effects of nutrient enrichment on arbuscular mycorrhizal fungi, plant diversity and ecosystem multi‐functionality. - New Phytologist. |
|  | [3] Fry, E. L. et al. 2018. Soil multifunctionality and drought resistance are determined by plant structural traits in restoring grassland. - Ecology 99: 2260–2271. |
|  | [4] Erktan, A. et al. 2018. Two dimensions define the variation of fine root traits across plant communities under the joint influence of ecological succession and annual mowing. - J. Ecol. 106: 2031–2042. |
|  | [5] Prieto, I. et al. 2015. Root functional parameters along a land-use gradient: evidence of a community-level economics spectrum. - J. Ecol. 103: 361–373. |
|  | [6] Kleinebecker, T. et al. 2014. Evidence from the real world: 15 N natural abundances reveal enhanced nitrogen use at high plant diversity in Central European grasslands. - J. Ecol. 102: 456–465. |
|  | [7] Craine, J. M. et al. 2001. The relationships among root and leaf traits of 76 grassland species and relative abundance along fertility and disturbance gradients. - Oikos 93: 274–285. |
|  | [8] Ostonen, I. et al. 2007. Specific root length as an indicator of environmental change. - Plant Biosystems - An International Journal Dealing with all Aspects of Plant Biology 141: 426–442. |
|  | [9] Reynolds, H. L. and D'Antonio, C. 1996. The ecological significance of plasticity in root weight ratio in response to nitrogen: Opinion. - Plant Soil 185: 75–97. |
|  | [10] Schenk, H. J. and Jackson, R. B. 2002. Rooting depths, lateral root spreads and below-ground/above-ground allometries of plants in water-limited ecosystems. - Journal of Ecology 90: 480–494. |
|  | [11] Giehl, R. F. and Wirén, N. von 2014. Root nutrient foraging. - Plant Physiology 166: 509–517. |
|  | [12] Reader, R. J. et al. 1993. A Comparative Study of Plasticity in Seedling Rooting Depth in Drying Soil. - J Ecol 81: 543. |
|  | [13] Klimešová, J. and Herben, T. 2015. Clonal and bud bank traits: patterns across temperate plant communities. - J Veg Sci 26: 243–253. |
|  | [14] Jones, J. B. 2017. Plant nutrition and soil fertility manual, second edition. - CRC Press. |
|  | [15] Treseder, K. K. 2004. A meta-analysis of mycorrhizal responses to nitrogen, phosphorus, and atmospheric CO2 in field studies. - New Phytologist 164: 347–355. |
|  | [16] Zhang, X. et al. 2019. Effects of precipitation change on fine root morphology and dynamics at a global scale: a meta-analysis. - Canadian Journal of Soil Science 99: 1–11. |

**Appendix S5** Percentage of variance explained and eigenvalues (printed inside the bars) for the four principal component analyses we did: (A) Above-belowground; (B) Belowground; (C) Above-belowground with plant functional type; (D) Belowground with plant functional type. The two first PCs explain about 55-60% of the total variance in each analysis.
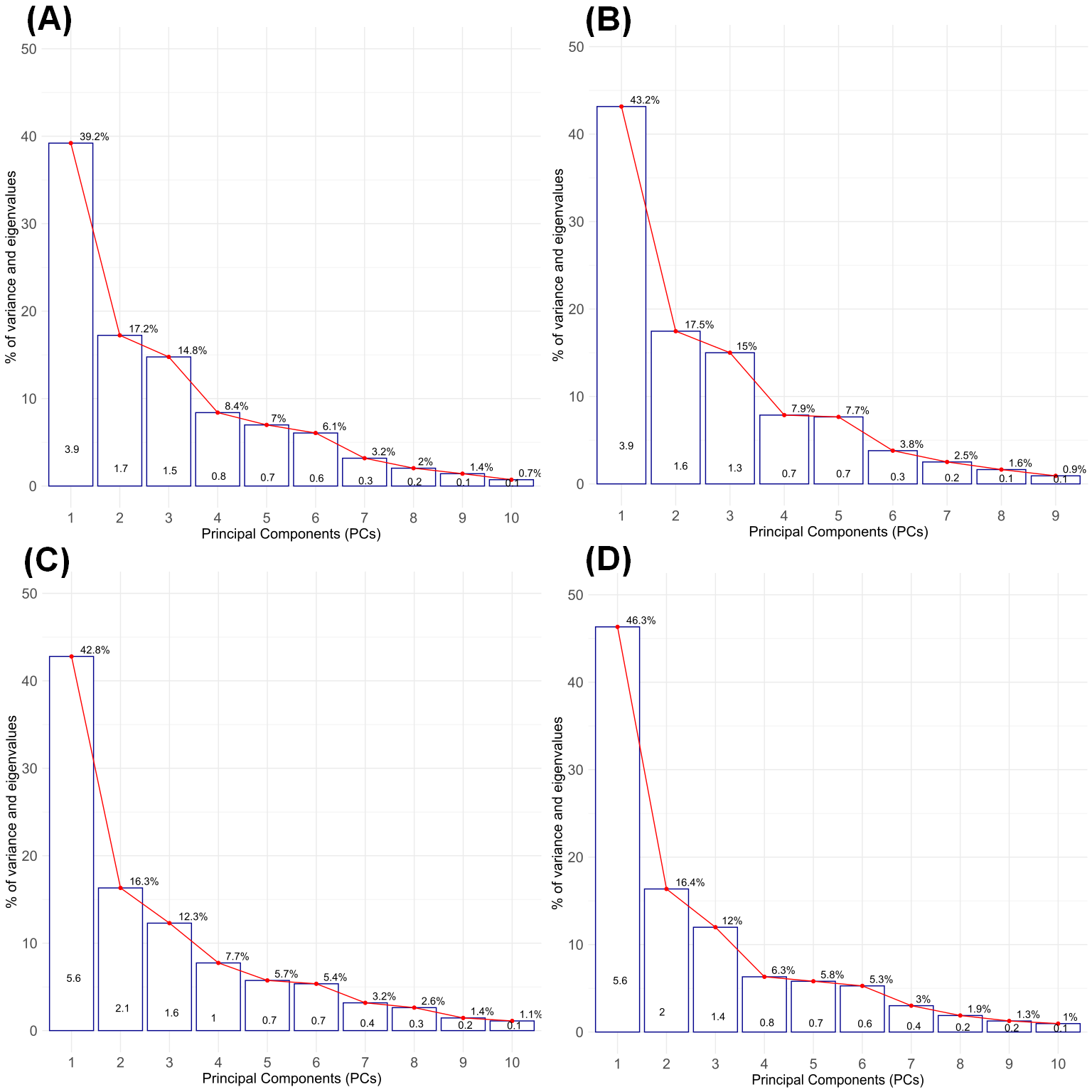

**Appendix S6** Two-dimensional projections of the two first PCs resulting from the four PCAs on (A) Above-belowground (also shown in Fig. 2); (B) Belowground; (C) Above-belowground with plant functional type; (D) Belowground with plant functional type. The two first PCs explain about 55% of the total variance in each configuration. _CWM_*Specific leaf area*, as the sole leaf trait is shown in green. The three plant functional types Poales, forbs (non-Fabaceae dicotyledons) and Fabaceae are shown in red. The scores of the 150 grassland plots used for the PCA are shown in red (Alb region, N=50), brown (Hainich region, N=50) and blue (Schorfheide region, N=50). The first PC (‘Collaboration’) is more characterized by CWMs of traits related to the ‘collaboration’ gradient. It associates thick fine roots with mycorrhizal colonization (‘outsourcing’), opposed to highly branched roots with high specific root length (‘do-it-yourself'). High proportion of Poales are characteristic of the 'do-it-yourself' strategy. The PC2 (‘Conservation’) is more strongly characterized by CWMs of traits related to the plant economic spectrum and vegetative regeneration potential. It associates high root weight ratio with large bud banks and high root tissue density (‘slow’), opposed to high specific leaf area (‘fast’). High proportions of forbs and Fabaceae are characteristic of the ‘outsourcing’ strategy. A high proportion of Fabaceae is also associated with ‘fast’, deep rooting communities, whereas a high proportion of forbs is associated with more ’slow’, regenerative communities.

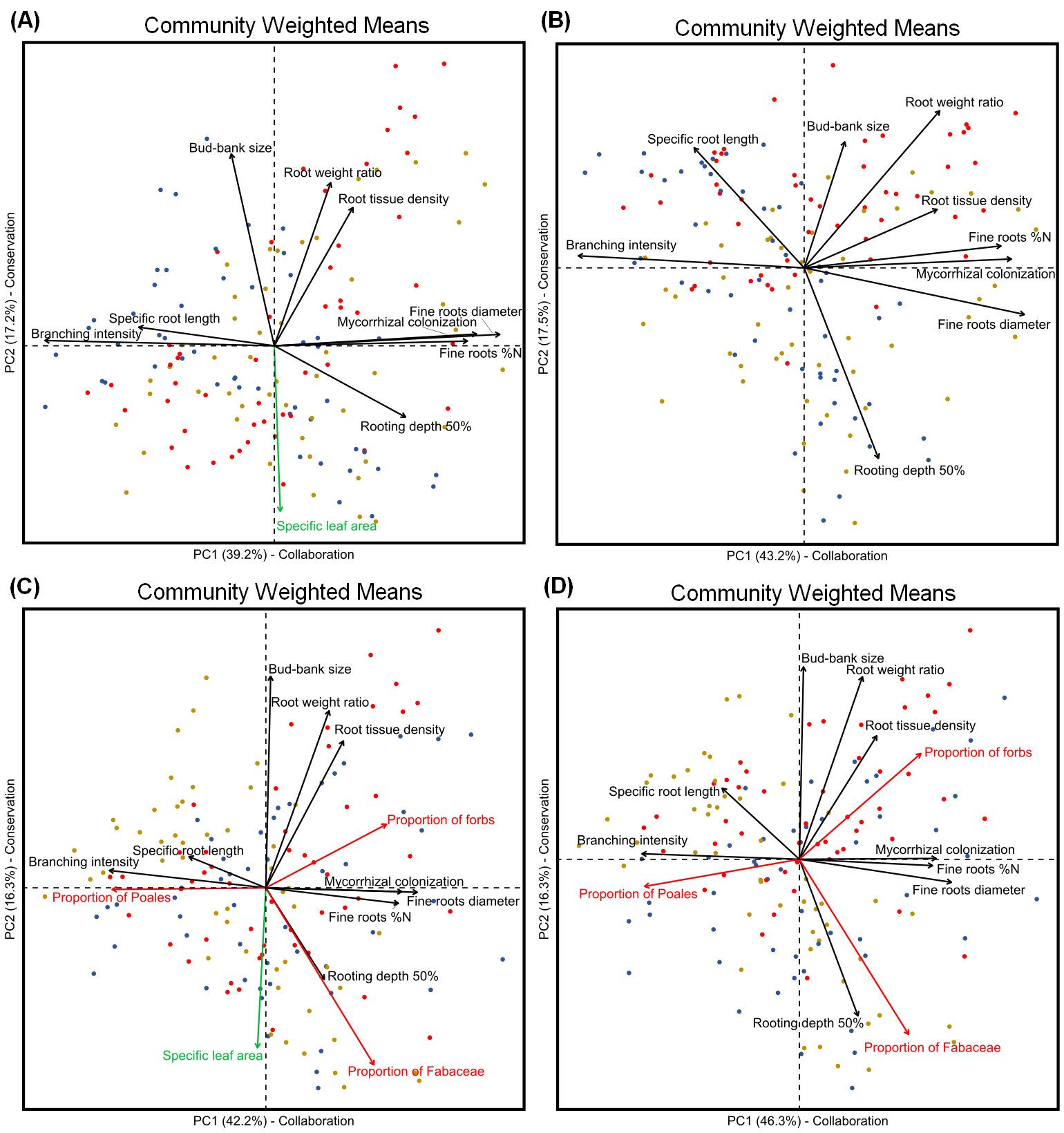

**Appendix S7** Two-dimensional projections of the two first PCs resulting from the four PCAs on (A) Above-belowground with the entire set of species included for CWMs calculations (also shown in Fig. 2); (B) Above-belowground excluding non-Fabaceae forbs for CWMs calculations; (C) Above-belowground excluding Poales for CWMs calculations; (D) Above-belowground excluding Fabaceae in CWMs calculations. _CWM_*Specific leaf area*, as the sole leaf trait is shown in green. The scores of the 150 grassland plots used for the PCA are shown in red (Alb region, N=50), brown (Hainich region, N=50) and blue (Schorfheide region, N=50). Removing Fabaceae or Poales for CWMs calculation has significant effects in modifying CWMs loadings on the two first PC, indicating that belowground traits relationships depend on plant functional types when considered at the community scale.

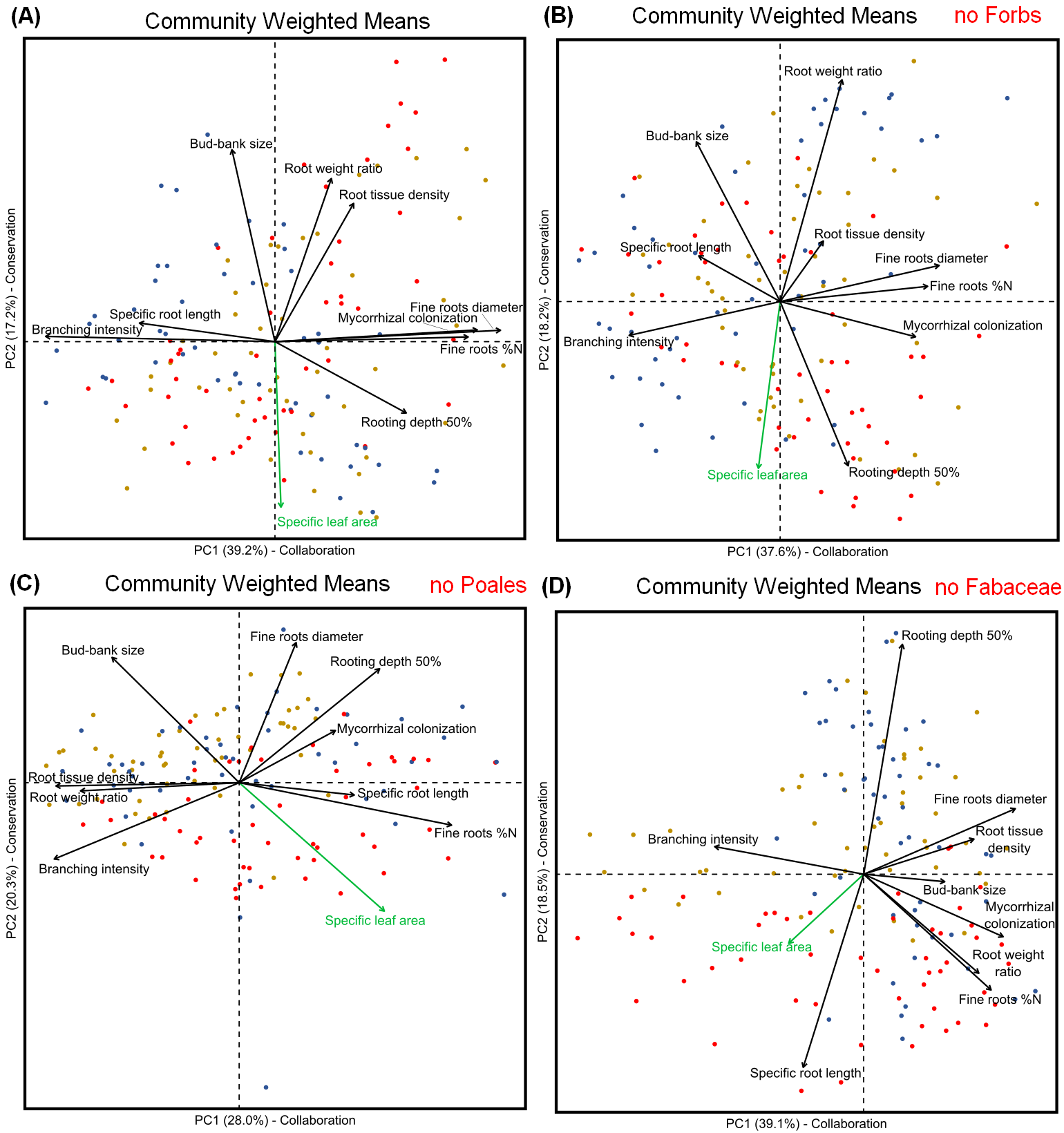

**Appendix S8** Estimates from linear models of environmental variables effects on the grassland plots functional identity, as captured by their scores for PC1 - Collaboration and PC2 - Conservation-regeneration, of the four PCAs on (A) Above-belowground (also shown in Fig. 2); (B) Belowground (corresponding to Appendix S6b); (C) Above-belowground with plant functional type (corresponding to Appendix S6c); (D) Belowground with plant functional type (corresponding to Appendix S6d). On the y-axis are the 9 environmental variables retained as predictors. The error bars around the estimates are standard errors. Significant (* for p < 0.05 ; ** for p < 0.01 ; *** for p < 0.001 ) negative and positive estimates are marked in red and blue, respectively. Non-significant (p > 0.05) estimates are marked in grey. Marginally significant (p < 0.10) estimates are marked with †.

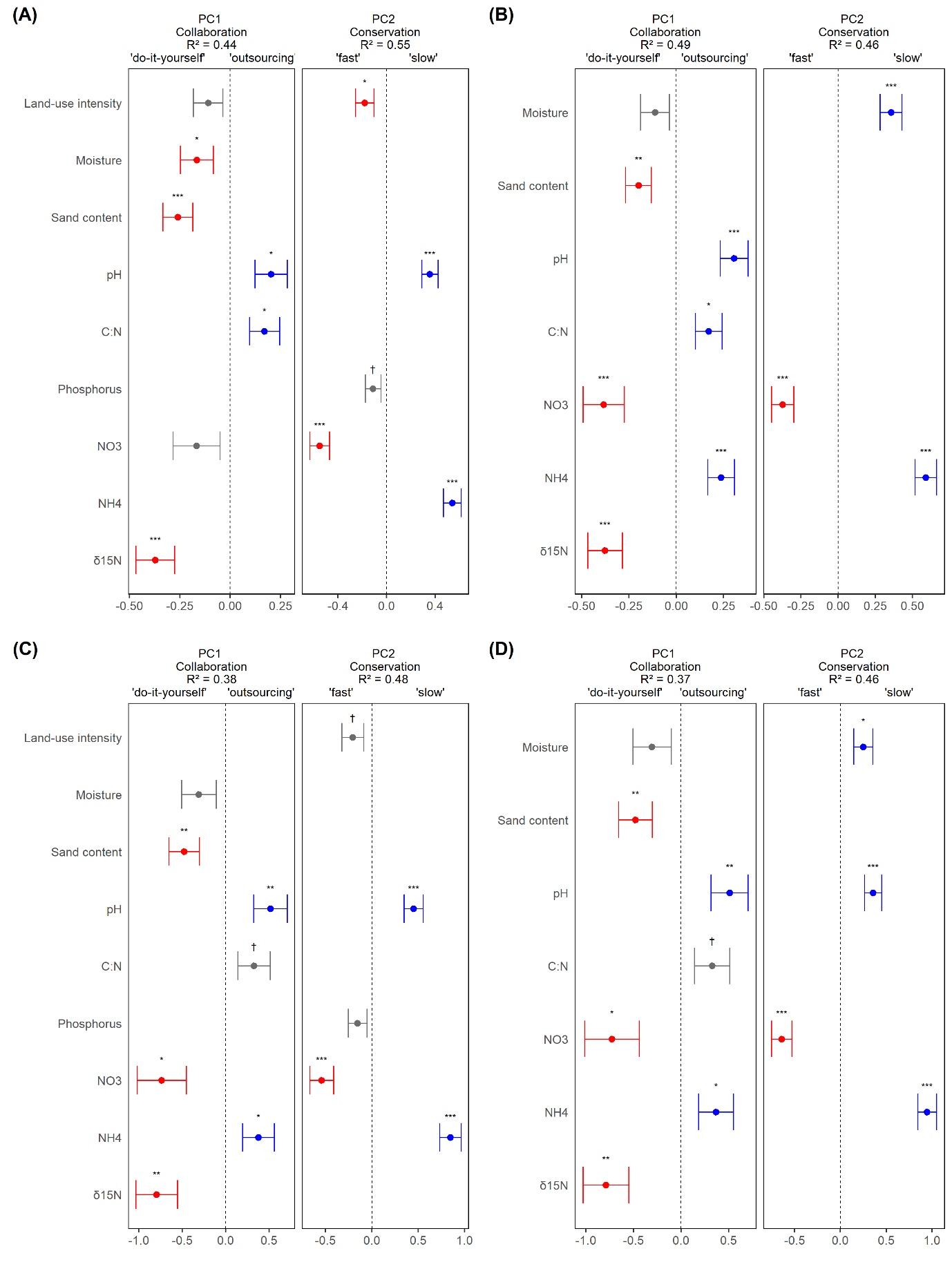

**Appendix S9** Loadings on the ten PCs of the Above-belowground PCA (as shown in Fig. 2 for PC1 and PC2 only). All the traits have strong loadings either on PC1 or PC2, representing 55-60% of the total variance. PC3 has a strong loading of _CWM_*Rooting depth 50%* and captures the differences in soil depth across the three regions (see Appendix S2 and Appendix S13). _‘_Oblimin’ rotation was used on PC1 and PC2 to accentuate the loading of CWMs on each PC.

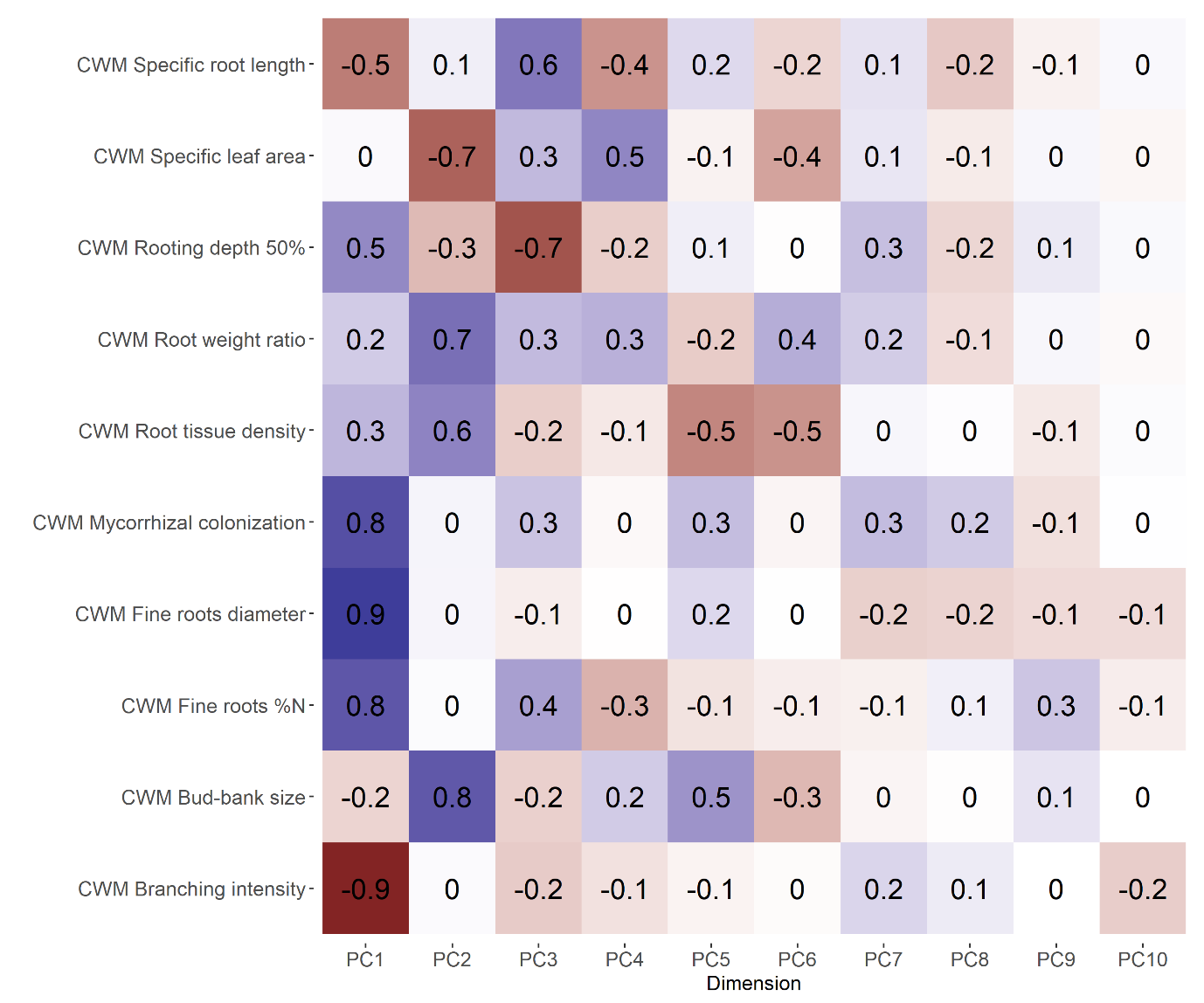

**Appendix S10** Correlation matrix of community weighted means (CWMs), proportions of three plant functional types, and PC scores for the two first PCs of the Above-Belowground PCA (as shown in Fig. 2). Pearson’s correlation coefficients are displayed. Negative correlations are in red, positive correlations are in blue, and non-significant correlations are indicated with a cross. Forbs = non-Fabaceae dicots. Multiannuals = biennials plus perennials.

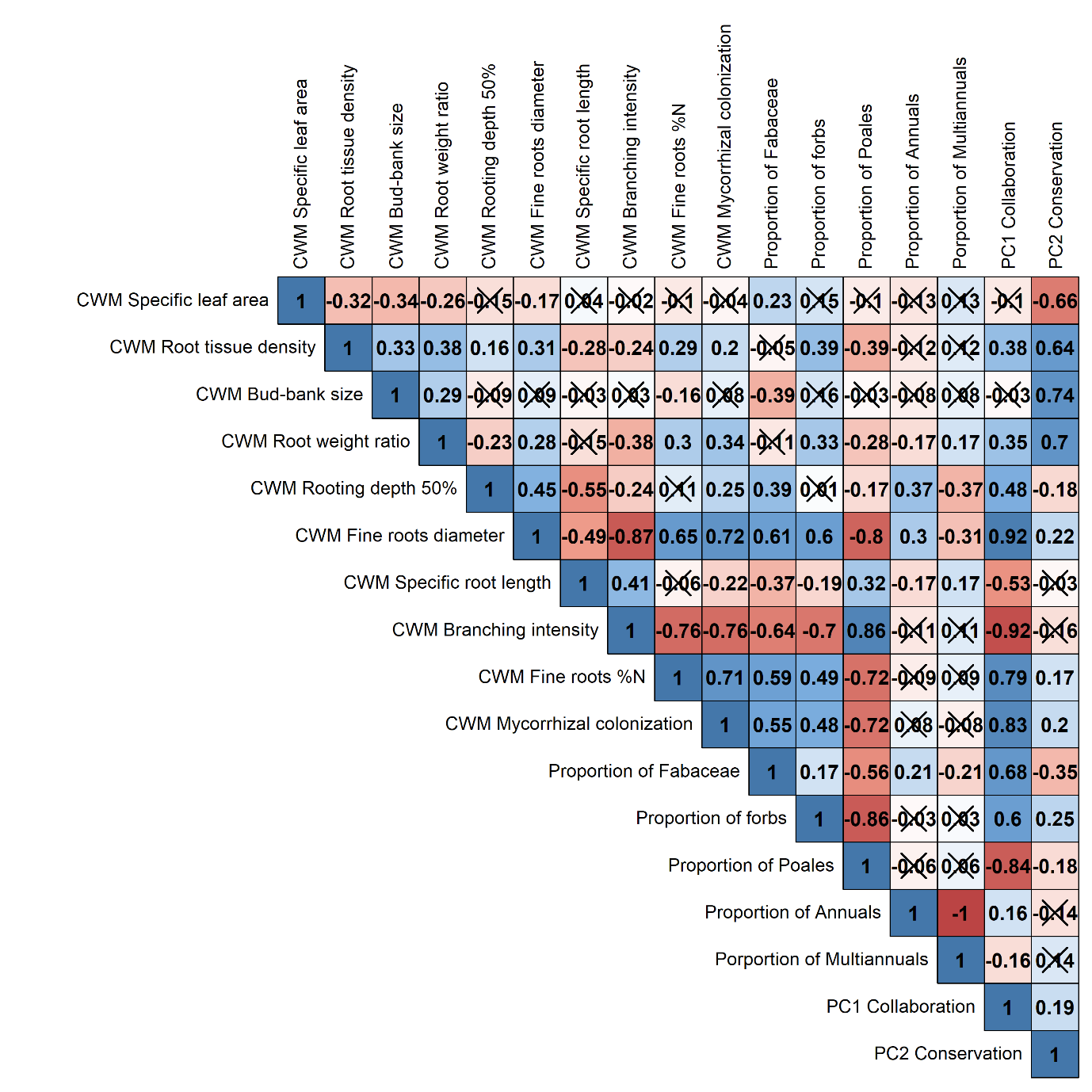

**Appendix S11** Correlation matrix of the environmental variables and proportions of three plant functional types. Bulk density wasn’t used directly as a predictor in models because of its strong correlation with several other soil parameters, resulting in large variance inflation factors (VIF), and was used indirectly to express soil parameters as unit of volume instead of mass when possible. Soil depth wasn’t used in models because of its insufficient sampling (N=144 and limited to 100 cm sampling, whereas for all the other predictors, N=150 and sampling values were not censored). Pearson’s correlation coefficients are displayed. Negative correlations are in red, positive correlations are in blue, and non-significant correlations are indicated with a cross.

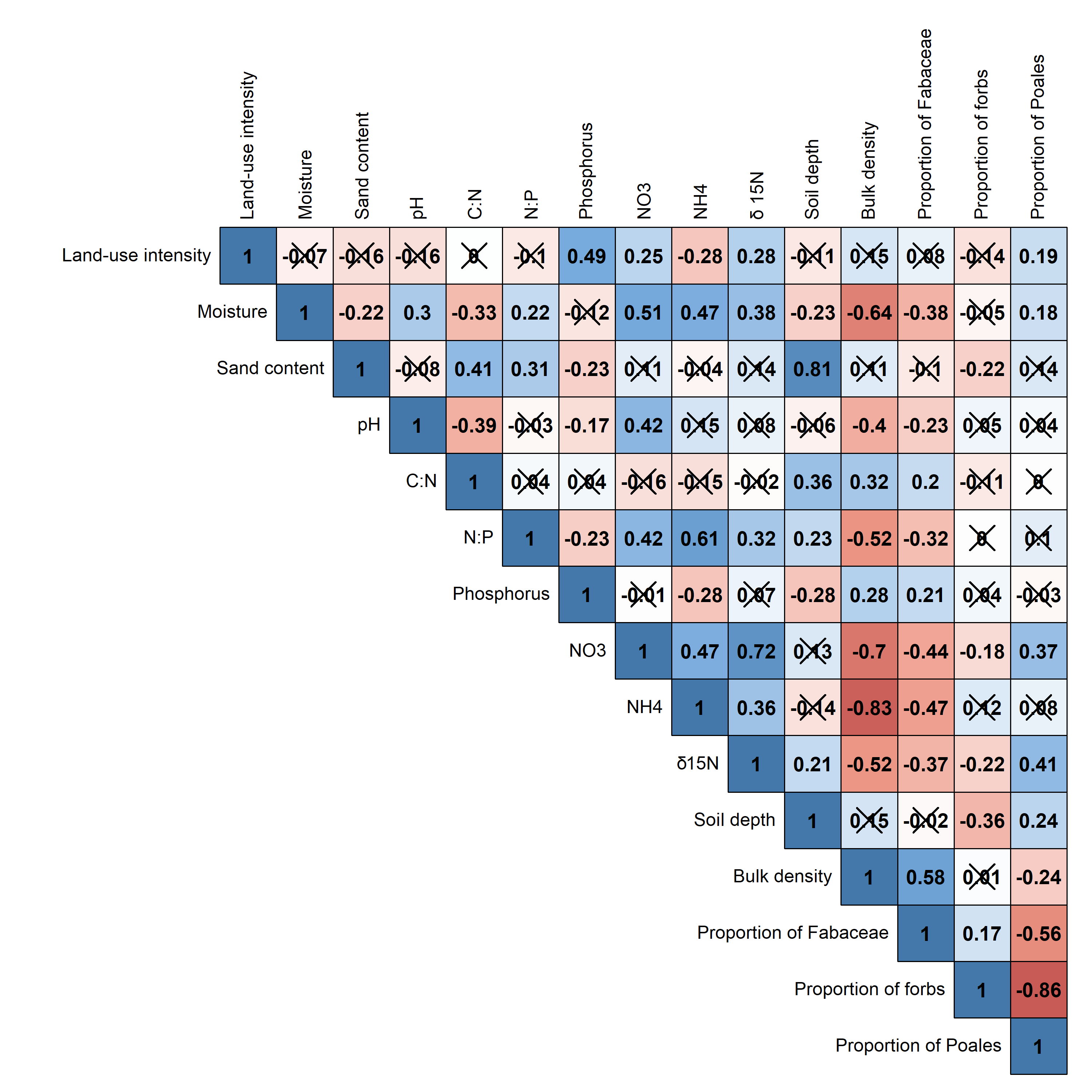

**Appendix S12** Estimates from linear models of environmental variables effects on the proportion of plant functional types. On the y-axis are the 9 environmental variables and the two regions retained by the models as predictors. The error bars around the estimates are standard errors. Significant (* for p < 0.05 ; ** for p < 0.01 ; *** for p < 0.001 ) negative and positive estimates are marked in red and blue, respectively. Non-significant (p > 0.05) estimates are marked in grey. Marginally significant (p < 0.10) estimates are marked with †. Forbs = non-Fabaceae dicots.

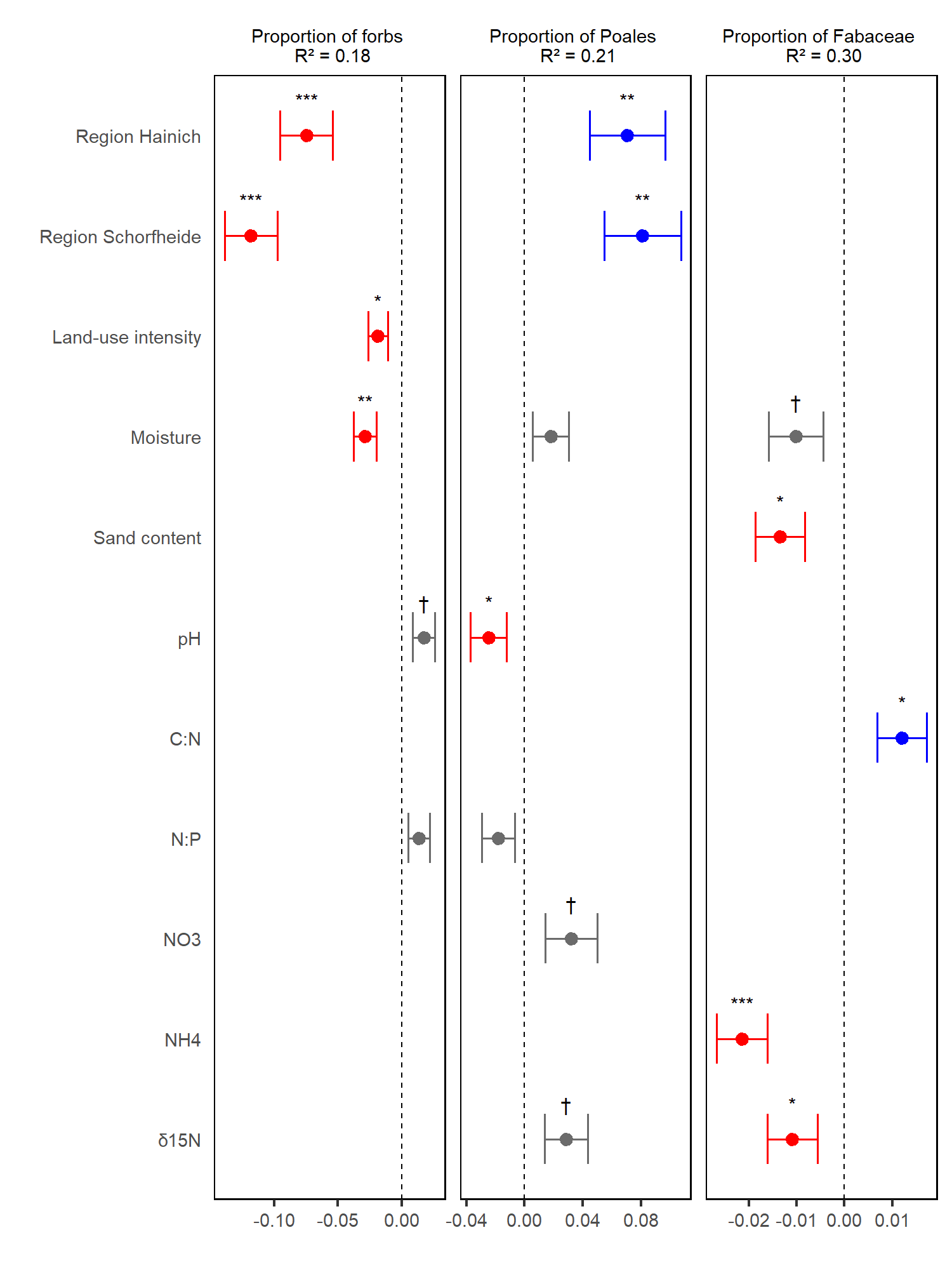

**Appendix S13** Estimates from linear models of environmental variables effects on the scores for the third to sixth PC of the Above-belowground PCA (as in Fig. 3 but for subsequent PC). Combined with PC1 and PC2, they together explain more than 90% of the total variance (see Appendix S5). On the y-axis are the 7 environmental variables and the two regions retained by the models as predictors. The error bars around the estimates are standard errors. Significant (* for p < 0.05 ; ** for p < 0.01 ; *** for p < 0.001 ) negative and positive estimates are marked in red and blue, respectively. Non-significant (p > 0.05) estimates are marked in grey. Marginally significant (p < 0.10) estimates are marked with †.

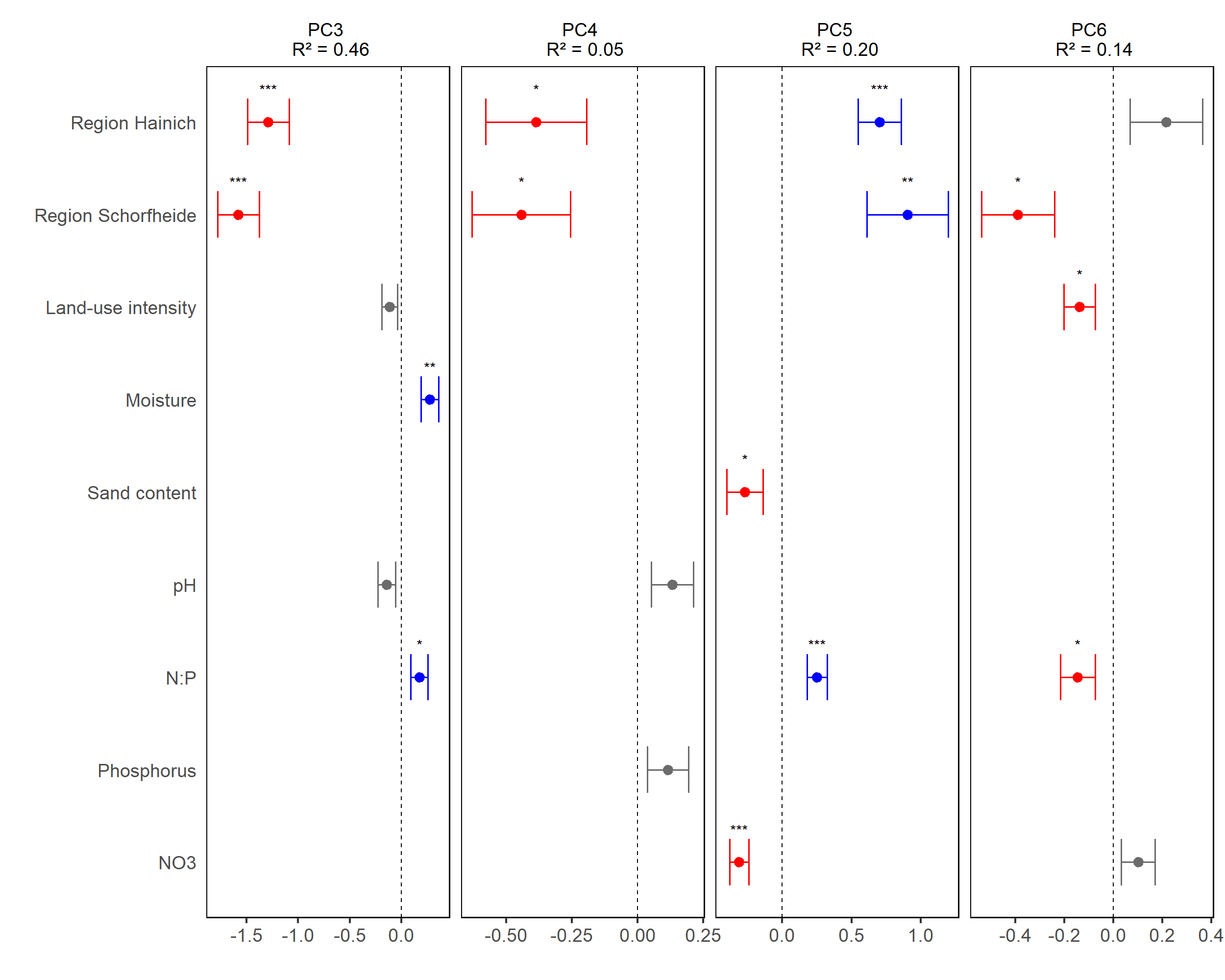

**Appendix S14** Estimates from linear models of environmental variables effects on the grassland plots community weighted means (CWMs) for each of the ten traits used in the four PCAs. On the y-axis are the environmental variables and regions retained by the models as predictors. The error bars around the estimates are standard errors. Significant (* for p < 0.05 ; ** for p < 0.01 ; *** for p < 0.001 ) negative and positive estimates are marked in red and blue, respectively. Non-significant (p > 0.05) estimates are marked in grey. Marginally significant (p < 0.10) estimates are marked with †.

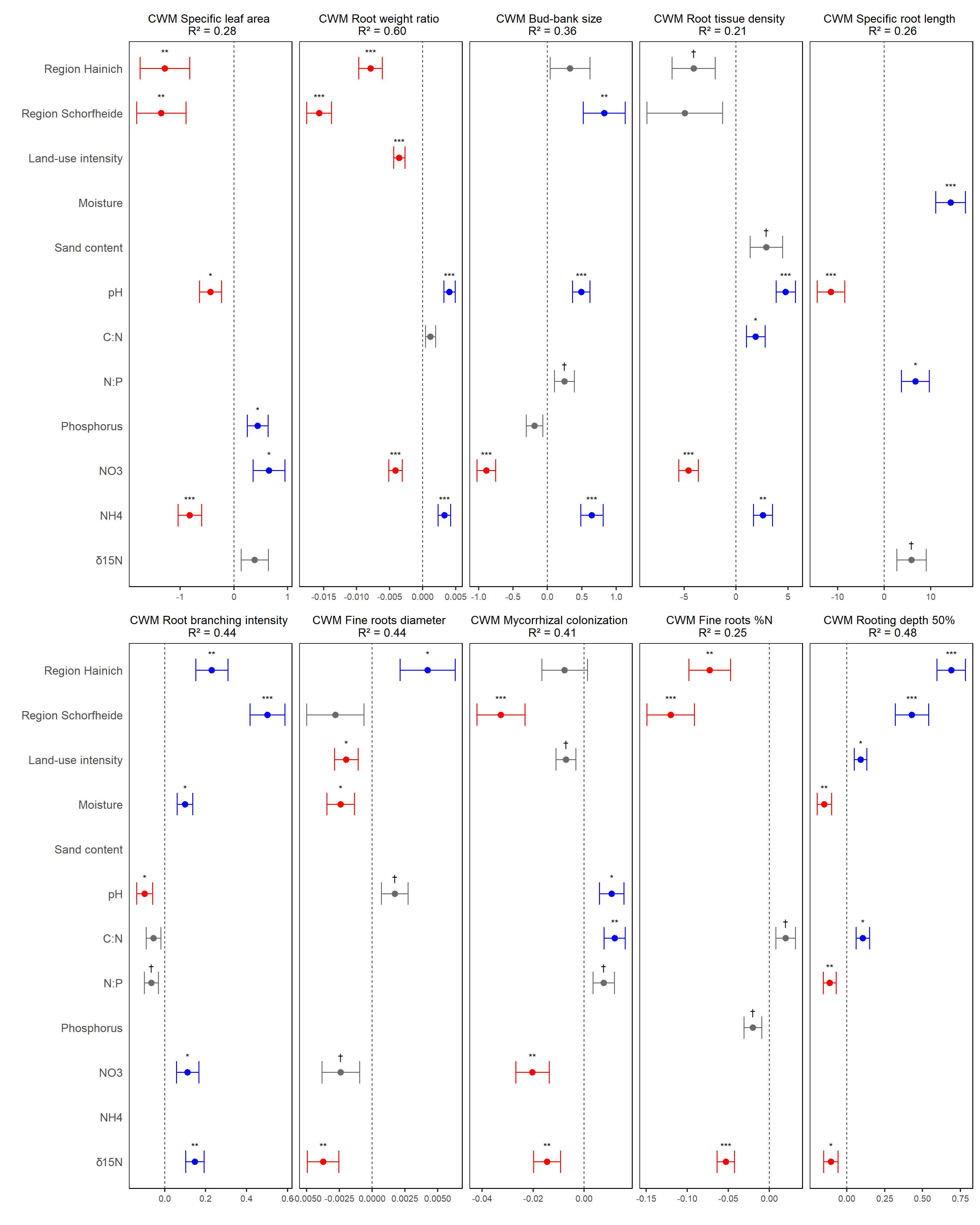

**Appendix S15** Indicator species or taxa for combinations of the dimensions of CWMs

*Methods*

To identify which species or taxa were associated with particular combinations of the dimensions of the CWMs of traits, we subsetted the grassland plots based on their scores on PC1 and PC2 of the Above-Belowground PCA (Fig. 2) into four quadrants, corresponding to the four combinations of positive and negative coordinates of the two PCs. For each quadrant, we calculated the percentage of cover represented by the ten most common species or taxa in the plots. We then calculated Pearson’s phi coefficient of association between species or taxa and the four PCA quadrants, used as sites (Cáceres & Legendre, 2009), with the function multipatt() from the package ‘indicspecies’ (Cáceres & Jansen, 2015).

*Summary diagram of the two main dimensions of the CWMs of belowground traits in German grasslands and their variation along environmental gradients.*

PC1 of the “Above-belowground PCA” corresponds to a ‘conservation’ gradient (vertical) with ‘fast’ and ‘slow’ strategies. PC2 corresponds to a ‘collaboration’ gradient (horizontal) with ‘do-it-yourself’ and ‘outsourcing’ strategies. Therefore, communities can be classified into four categories, according to their position in one of the four quadrants of the projection system of PC1 and PC2. The size of the quadrants reflects the number of plots in each quadrant ('Slow & Do-it-yourself': N = 28 plots; 'Slow & Outsourcing': N = 37 plots; 'Fast & Do-it-yourself': N = 46 plots; 'Fast & Outsourcing': N = 39 plots. Inside each quadrant, we indicate the ten dominant species or taxa with the highest cover in the plots. They contributed from 40.1% of the total cover for 'Slow & Outsourcing' to 66% for 'Fast & Outsourcing'. Highlighted in different colors are the indicator species or taxa for one or several quadrants. Red = one quadrant; Green = two quadrants; Blue = three quadrants. The largely overlapping list of dominant species or taxa indicates that variation in CWMs relies on the turnover of the same set of dominant species. It is worth mentioning that although Poales are overall characterized by lower mycorrhizal colonization than forbs, grasses with high mycorrhizal colonization are dominant species in the ‘Slow & Outsourcing‘ quadrant. This indicates that the gradient from ‘Do-it-yourself’ to ‘Outsourcing’ communities is not only explained by replacement of Poales by forbs, but also by changes within the Poales group itself. The barplots are highlighting, for two traits characteristic of each quadrant, their difference between their (scaled) CWMs and the CWMs for all communities. At the four extremities of the two-dimensional space, we highlighted which environmental gradients are associated with which kind of communities, and the font size is scaled with the coefficients from regression models as shown in Fig. 3.

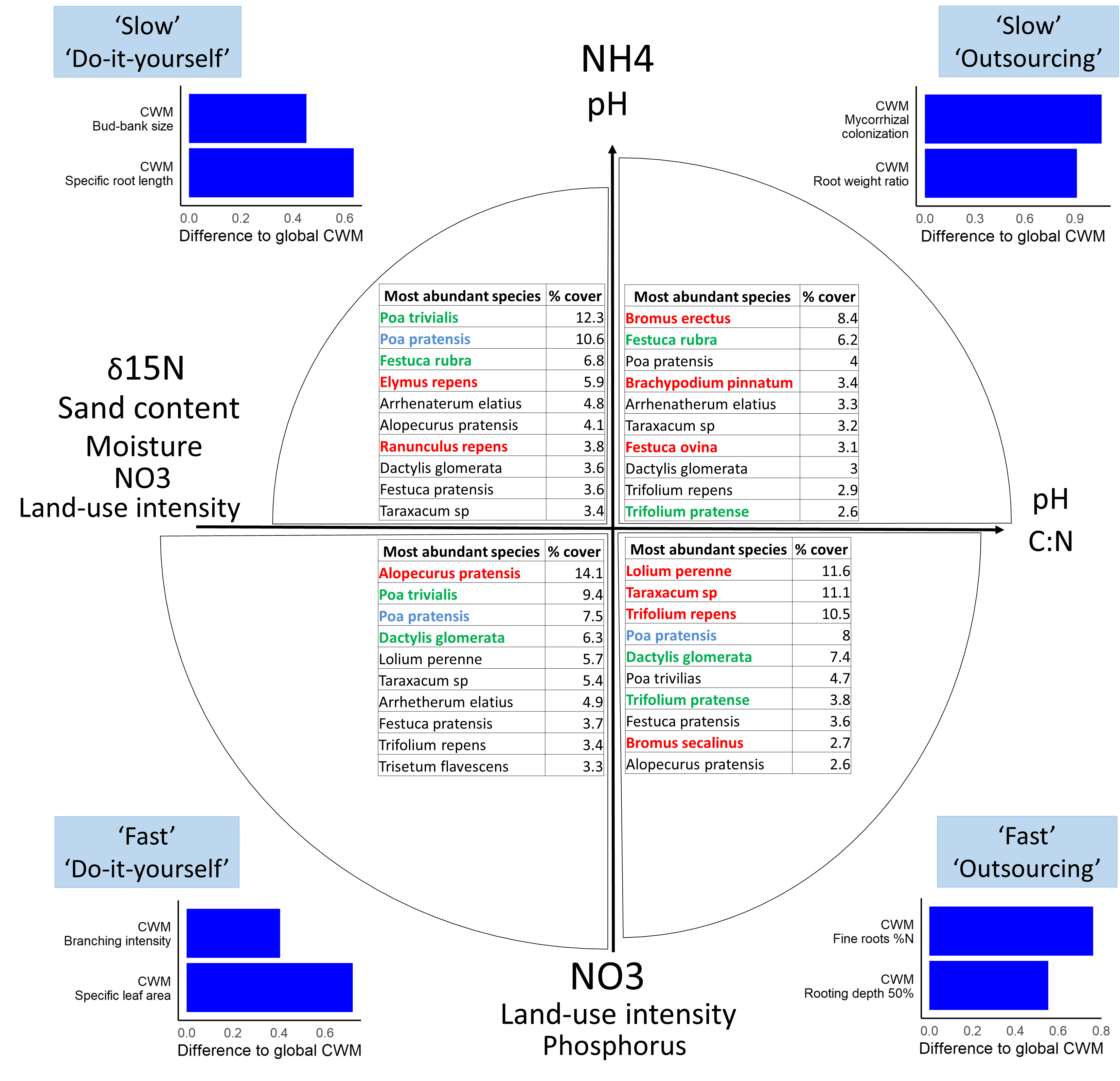

**Appendix S16** (A) The two first PCs of the Above-belowground PCA at the species level (instead of the community level), explaining 40.2% of the total variance in species-level trait means. The sole aboveground trait that we included, specific leaf area, is shown in green. The scores of the 218 species, covered by the morphology experiment are shown as dots, and imputation by the mean are used for the other traits to fill the gaps between their sample size and the sample size of the morphology experiment for the following number of species (specific leaf area: N=21; fine roots %N: N=20; mycorrhizal colonization: N=143; bud-bank size: N=7, rooting depth 50%: N=39). The plant functional types of the species are indicated in blue: Fabaceae (N=23); red: non-Fabaceae forbs (N=151); brown: Poales (N=44). (B) Pearson’s correlation coefficients between the traits at the species level. Negative correlations are in red, positive correlations are in blue, and non-significant correlations are indicated with a cross.

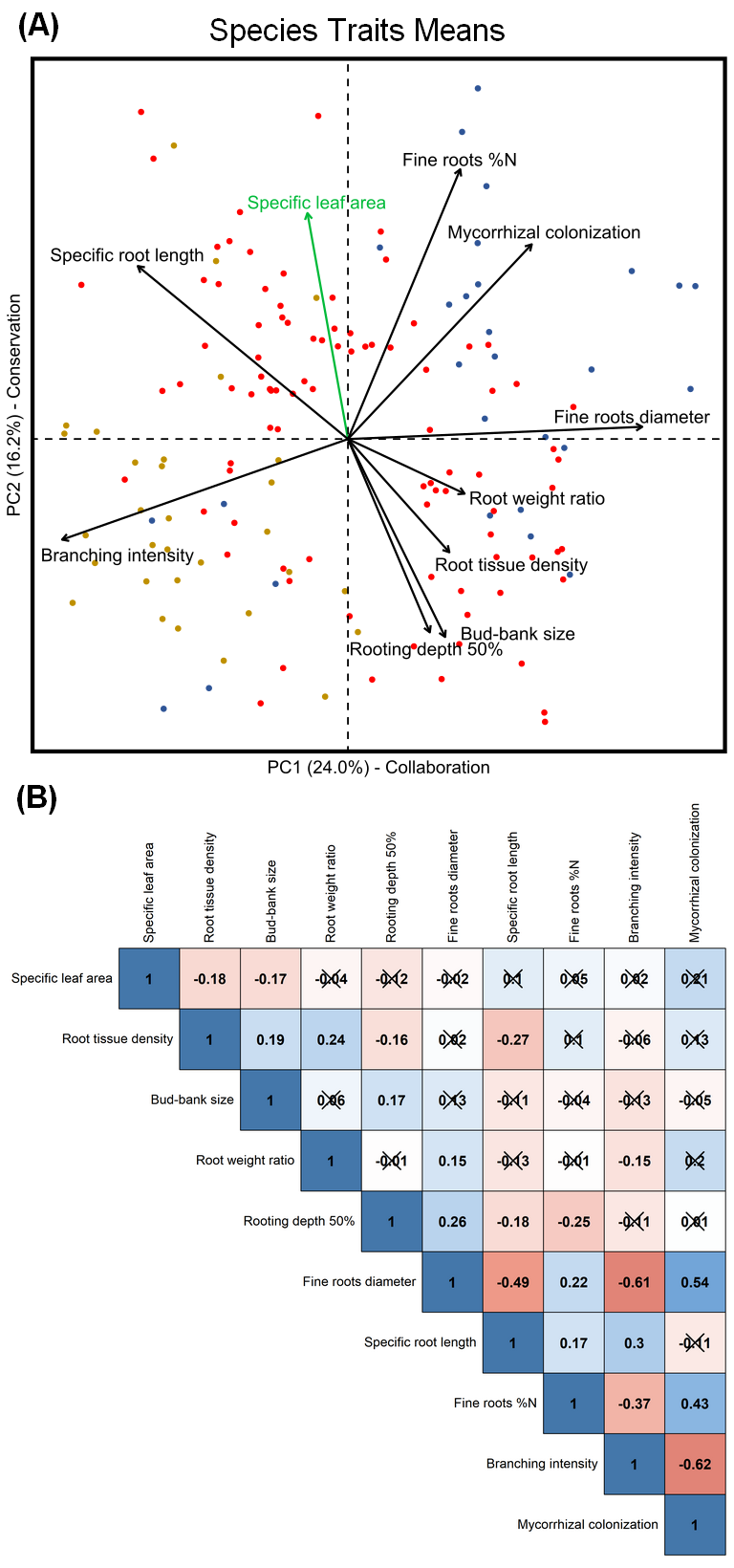
